# STEPS 4.0: Fast and memory-efficient molecular simulations of neurons at the nanoscale

**DOI:** 10.1101/2022.03.28.485880

**Authors:** Weiliang Chen, Tristan Carel, Omar Awile, Nicola Cantarutti, Giacomo Castiglioni, Alessandro Cattabiani, Baudouin Del Marmol, Iain Hepburn, James G King, Christos Kotsalos, Pramod Kumbhar, Jules Lallouette, Samuel Melchior, Felix Schürmann, Erik De Schutter

## Abstract

Recent advances in computational neuroscience have demonstrated the usefulness and importance of stochastic, spatial reaction-diffusion simulations. However, ever increasing model complexity renders traditional serial solvers, as well as naive parallel implementations, inadequate. This paper introduces a new generation of the STochastic Engine for Pathway Simulation (STEPS) project, denominated STEPS 4.0, and its core components which have been designed for improved scalability, performance, and memory efficiency. STEPS 4.0 aims to enable novel scientific studies of macroscopic systems such as whole cells while capturing their nanoscale details. This class of models is out of reach for serial solvers due to the vast quantity of computation in such detailed models, and also out of reach for naive parallel solvers due to the large memory footprint. Based on a distributed mesh solution, we introduce a new parallel stochastic reaction-diffusion solver and a deterministic membrane potential solver in STEPS 4.0. The distributed mesh, together with improved data layout and algorithm designs, significantly reduces the memory footprint of parallel simulations in STEPS 4.0. This enables massively parallel simulations on modern HPC clusters and overcomes the limitations of the previous parallel STEPS implementation. Current and future improvements to the solver are not sustainable without proper software engineering practices. For this reason, we also give an overview of how the STEPS codebase and the development environment have been updated to follow modern software development practices. We benchmark performance improvement and memory footprint on three published models with different complexities, from a simple spatial stochastic reaction-diffusion model, to a more complex one that is coupled to a deterministic membrane potential solver to simulate the calcium burst activity of a Purkinje neuron. Simulation results of these models suggest that the new solution dramatically reduces the per-core memory consumption by more than a factor of 30, while maintaining similar or better performance and scalability.

## 1 Introduction

For several decades computational modeling has progressively proven its importance in the frontiers of neuroscience research, covering a wide range of research domains and disciplines: from sub-cellular molecular reaction-diffusion dynamics to whole-brain neural network simulations. Breakthroughs in experimental methods and community-driven data sharing portals have significantly increased the amount of available experimental data, enabling the advance of complex data-driven modeling and analysis. These efforts are further enhanced by large collaborative projects such as the US BRAIN initiative (Insel et al., 2013), and the EU Human Brain Project, (Markram et al., 2011), where complex computational modeling plays an essential role. The rapid progress of neuroscience modeling brings critical advances to our understanding of neuronal systems, yet unprecedented challenges to simulator software development have emerged from two primary directions: first, the need to simulate convoluted neuronal functionalities across multiple spatio-temporal scales, and second, the requirement of simulating such systems with extraordinary efficiency.

### 1.1 The STEPS project and its applications

The STochastic Engine for Pathway Simulation (STEPS) project has also evolved following the above trends over the years. The STEPS project started as a mesoscopic scale stochastic reaction-diffusion solution (Hepburn et al., 2012) driven by a spatial variant of the renowned Gillespie SSA method (Gillespie, 1977). Over the years, serial STEPS has contributed to a wide range of research domains, such as studies on viral RNA degradation and diffusion (Schelker et al., 2016), longitudinal anomalous diffusion in neuron dendrites (Mohapatra et al., 2016), long-term depression in cerebellar Purkinje cells (Zamora Chimal and De Schutter, 2018), and calcium signaling in astrocytic processes (Denizot et al., 2019). We gradually expanded STEPS to support electrical potential calculation on tetrahedral meshes with the EField solver (Hepburn et al., 2013), allowing combined simulation of reaction-diffusion and membrane potential dynamics on a single mesh reconstruction of neuronal morphology. This solution was important for the research of the stochastic effect on dendritic calcium spiking variability (Anwar et al., 2013). However, it was soon clear to us that the serial nature of STEPS was the major bottleneck for simulating such complicated models; even a sub-branch of a Purkinje neuron often took weeks to complete one realization. This issue was partially addressed in STEPS 3.0 by introducing the parallel operator splitting method to the reaction-diffusion solution (Hepburn et al., 2016; Chen and De Schutter, 2017), which aided research such as platform development for automatic cancer treatment discovery (Stillman et al., 2021). A parallel EField implementation supported by the PETSc library (Abhyankar et al., 2018) was added to STEPS 3.1. The parallel solution dramatically improved performance by thousand folds compared to the serial counterpart, making it possible to model a complete neuron with detailed morphology and channel mechanisms (Chen et al., in press).

### 1.2 The need of a new parallel solver

Moving to parallel STEPS has greatly improved performance compared to the serial solution. However, as the hardware and software of high-performance computing have advanced in recent years, noticeable bottlenecks have been observed in modeling applications with STEPS. The main objective of this article is to identify these bottlenecks and address them with a new parallel implementation.

For many scientific applications, the memory capacity of High-Performance Computing (HPC) systems is one of the main constraints for running simulations at scale. A large number of today’s HPC systems have about 2∼3GB of main memory per core (Zivanovic et al., 2017). This is an improvement compared to previous BlueGene-like systems where memory capacity is typically ∼1GB per core. Current systems are increasingly heterogeneous with the use of accelerators such as GPUs. The memory capacity of such a system is significantly lower compared to what is commonly available on host CPUs. We have seen a similar situation in the case of Intel Knights Landing (KNL) processors (Sodani et al., 2016), with a total capacity of approximately 0.2GB per core. The next generation of processors such as Intel Sapphire Rapids (SPR) will most likely have a similar per-core memory capacity. This poses a significant challenge to application developers: on the one hand the raw computing power is significantly increasing with architectures like GPUs, while on the other hand maintaining a low memory footprint becomes increasingly important to achieve better performance.

One major limitation of the existing parallel implementation in STEPS comes from the mesh data architecture inherited from the serial solution. While bridging the gap between serial and parallel STEPS and making many non-parallel components reusable, the serial nature of the design requires the complete data of the whole mesh and the molecule state of each mesh element to be stored in every computing core. This poses a hard limit on the maximum model size determined by the per-core memory availability, the model complexity, and the mesh size. Thanks to the support from the parallel solver, realistic simulations with a large number of chemical reactions for a great period of biological time can now be accomplished in a reasonable computing time. However, this in turn raises research interests in even more complicated models and more realistic morphologies, reaching the limits of the implementation. The memory constraints in modern HPC systems further amplify such limitations.

The solution to this issue is a new parallel implementation constructed on the foundation of a sophisticated distributed mesh library, Omega_h (Ibanez and Roberts, 2018). Thanks to the distributed nature of the mesh library and the redesigns of other STEPS components, our new implementation dramatically reduces the memory footprint of the simulation while maintaining similar or better performance and scalability. This enables super-large scale simulations that were impossible with previous implementations due to hardware constraints.

### 1.3 Naming conventions and the structure of the article

To avoid confusion, we will hereby call the non-parallel, spatial STEPS solver “serial STEPS”, the existing parallel implementation reported in Chen and De Schutter (2017) as “STEPS 3”, and the new parallel implementation supported by Omega_h that we introduce in this paper as “STEPS 4”. Note that serial STEPS, STEPS 3 and STEPS 4 are all integrated solutions of the STEPS 4.0 release, and the users are free to choose any of them for their simulations based on the research requirements.

In the Methods section, we first describe our design principles and the implementation details of STEPS 4, and then introduce some software engineering techniques applied to the overall STEPS project for maintainability and efficiency improvements, which may be interesting to colleagues in the fields of computational neuroscience modeling or scientific software development. In the Results section, we present the validations of the implementation with a series of well-established models, followed by performance and scalability analysis of results. In Discussion, we further discuss the achievements, limitations and potential solutions of this study, as well as the future development plans for STEPS 4 and the STEPS project in general.

## 2 Methods

The STEPS development project follows three major methodological principles. First, it aims towards the researchers. A successful simulator should provide a friendly modeling interface and significantly reduces the need for manual coding efforts. Second, it focuses on improving performance. Because fundamentally, whether a simulator is a suitable option for a research project greatly depends on if the simulations can complete within the expected research time frame. Third, it aims to be future-proof. Since the first public release, the STEPS project has more than ten years of history. Over the years, many new standards and solutions in programming and software engineering have been established and become the new standard in software development. Some of them have been adopted in previous STEPS development, but more work is still required to ensure that the software development infrastructure is ready for future project expansions. The following sections detail how these principles are practically applied in the project.

### 2.1 API improvements for a user-friendly modeling environment

The field of computational neuroscience is recognized as very multi-disciplinary in nature, bringing researchers together from a wide variety of backgrounds such as biology, computer science and physics, and so STEPS has always been designed with a focus on supporting researchers from different backgrounds and with different levels of computational expertise. In addition, designing, running, and analyzing a spatial molecular model is a complex task that is far better suited to a scripting language such as Python as opposed to a standalone C++ application. Although it is known that such detailed molecular models are computationally expensive, it is important for the user that the software is fast and efficient, which is best achieved with a C++ backend. So very early on in STEPS’ development, it was decided to combine a scripting interface in Python through which the user interacts with the software and can build detailed spatial models, but with computations carried out in C++ for speed and efficiency (Wils and De Schutter, 2009). Of the scripting languages, Python was chosen due to its maturity and increasing user-base among the computational biology community, with many software packages in the field now providing a Python interface.

When designing an Application Programming Interface (API), software developers frequently must deal with trade-offs between efficiency, flexibility, and complexity. In such cases, decisions should be made based on the user expectations of the application. Since its first release, STEPS provides a Python-based API that covers extensive details of the entire simulation space with different resolutions. This gives the users extremely flexible control over the simulation according to their research needs, although it also increases the complexity of modeling and sometimes also with the cost of simulation efficiency. One good example is the task of acquiring the count of a given molecule species in the simulation. In STEPS, this value can be returned as a single global count, similar to the result in a traditional biochemical simulation under well-mixed conditions. Alternatively, it can also be returned as a distribution over a collection of Regions Of Interest (ROIs) predefined by the modeler. Furthermore, it is possible to increase the spatial resolution up to elemental scale, i.e., iterate over all tetrahedral sub-volumes of the mesh and acquire the individual molecule counts of each sub-volume using elemental-based data access functions. However, every Python call to the C++ kernel incurs a substantial performance overhead, which can add up to a bottleneck if the call is iterated on a massive number of mesh elements. Such operations may be even more costly under STEPS 4 as global synchronization of data may be needed. In order to reduce the overhead, batch access functions were introduced to assist element-iterative data queries of this type.

In previous STEPS releases, the choices of the above functions were purely decided by the modeler, thus the same functionality may differ between models, and sometimes with significant differences in terms of performance. In STEPS 3.6, a new set of API, named API2, has been introduced. API2 provides encapsulations and abstractions of previous APIs, thus reducing the programming workload for STEPS users and allowing them to better focus on model development. In this work, we have further developed this new API to suit the distributed mesh nature in STEPS 4. At the same time, we ensure that Python scripts written for previous STEPS versions are still entirely compatible with the new release without any modification. Users are encouraged to use API2 for future model developments as the old API system will be progressively deprecated in future releases.

In order to use the new API system, users need to `import steps.interface` before any other STEPS python imports. There is no tool to fully convert a script, which uses the previous API, to the new one. While some parts of models like reaction and diffusion rule declarations could be converted automatically to some extent, data querying and saving is done in a substantially different way with the new API and thus does not lend itself well to automatic conversion. Most notably, in API2, users do not explicitly call specific data querying methods during the simulation. Instead, the data to be saved is specified beforehand and STEPS dynamically decides which internal methods should be called, depending on which methods are available with the solver being used. Saving to files is now also done by STEPS internally, allowing the user to only focus on which data needs to be saved instead of on how to save it. Finally, although it is beyond the scope of this paper, API2 introduces some concepts from rule-based modeling (Chylek et al., 2013) that allow the specification of multi-state complexes. For some biochemical models, this can considerably reduce the number of chemical reactions that the modeler needs to declare. As with the automatic saving changes, this new feature does not lend itself well to automatic conversion. Code examples about API2 and comparison to the original API are provided as supplementary material (S1).

### 2.2 Code modernization and future-proofing

Although STEPS introduced many new features and additions in the following years since its first release in 2012 the core coding components and style remained relatively unchanged. With this in mind, in this work we have implemented various far-reaching changes in STEPS in general and adopted modern software design principles to STEPS 4 in particular. All these changes have the aim to reduce bugs, improve maintainability and usability of the code and increase the performance of time-critical data structures and routines.

First, we have adopted the C++17 standard for STEPS. This allowed us to take advantage of modern programming language features, increasing code expressiveness and compactness through meta-programming techniques such as SFINAE (Substitution Failure Is Not An Error). We have also removed raw-pointers in favor of references and other safer data-passing and access strategies provided by the C++ standard and the GSL Guidelines Standard Library. Second, we have reduced code branching and indirections using meta-programming techniques, which streamline code execution. Third, when choosing container data structures we avoid C++ standard template library associative containers, which are known to be very inefficient in terms of memory management and performance. Instead, we have designed a new optimized data structure, named `flat-multimap`, which is designed to provide an associative container stored in contiguous arrays, increasing data locality and thus increasing performance.

With the increased complexity of a software, there is a growing concern about introducing bugs in the code that remain undetected. In the best case, these bugs will lead to crashes during runtime. In the worst case, they may silently introduce erroneous results and non-reproducible behavior. Although STEPS runs an extensive validation set to try and ensure this doesn’t happen, it is difficult to make sure that every base is covered by such efforts. In an attempt to address this issue at least partially, we introduced C++ vocabulary types meant to indicate the compiler the different entities used in a STEPS simulation (e.g. `species`, `membrane`, `channel`, `patch`, etc., but also `tetrahedron`, `triangle`, etc.). For instance, vocabulary types can improve functions like below:

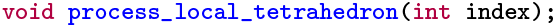

In the signature of this function, most of the information about the parameter is carried by the variable name and function name, which the compiler cannot use. For the compiler, `process_element` is only a function that takes a 32 bits integer in parameter. For the developer, this integer is an index of a tetrahedron, local to the current process. Vocabulary types allow us to transfer information traditionally held by the name of the symbols to the typing system by rewriting the signature of the function like this:

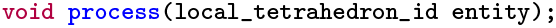

Thus, the compiler is now able to report an issue when the index of one type is erroneously passed to a function expecting another type.

Furthermore, by ensuring the code compiles with GCC, clang, AppleClang and Intel OneAPI, we ensure that language and system compatibility is maintained, further increasing code safety. Numerous compilation flags have been added into our build system, which allow us to spot and fix potential issues in the code early in the development process. We have also moved to a more module build design where features can be enabled via build configuration flags, which also benefits overall software architecture.

Last but not least, we have taken a number of steps to improve software sustainability beyond code modernization. To improve developer confidence and bug detection we have added continuous integration (CI) pipelines into the review process. Proposed patches are automatically built and tested before they can be merged into the development trunk. We have also created a STEPS package for the Spack (Gamblin et al., 2015) package manager. This not only adds a software distribution channel for HPC systems but also provides the developers with a comprehensive build environment that allows them to conveniently test STEPS with various dependency versions and build options. The choice of the underlying libraries (cf. section 2.3.2) plays an important role in ensuring that STEPS remains well maintainable, and easily extensible towards new features and use-cases while continuing to support the latest hardware architectures and parallel programming paradigms.

### 2.3 Implementing a parallel solver with distributed mesh backend

#### 2.3.1 Implementation criteria

To be able to make informed choices about the STEPS 4 implementation, we set early in the development a number of criteria by which to make decisions. Clearly the first and most important criterion is simulation runtime. The goal of the STEPS 4 implementation is to develop a new efficient solution for large-scale modeling with complex geometries. From a user’s perspective, the most straightforward and important concern is time-to-solution, how fast a simulation reaches a desired stopping time. For parallel simulations, another important concern is scalability. In high performance computing, parallel scalability is commonly described by two notions, strong scaling and weak scaling. The former describes runtime performance at increasing number of cores and a fixed problem size, while the latter scales the problem size with the number of cores. In practice, the problem size of a STEPS production simulation is often determined by the source materials. Thus, we focus on strong scaling as our parallel performance criterion. STEPS 4 is designed mainly for simulations that run on high performance computing clusters. As mentioned previously, one key characteristic of modern clusters is the large amount of computing cores together with the limited amount of per-core memory, thus memory footprint management is essential to support large scale simulations. We regard it as our third implementation criteria.

These criteria often affect each other in a simulation. For instance, the reduction of memory footprint could substantially improve the efficiency of memory caching, and further improve scalability. Therefore, we do not focus on individual criterion, but consider them as a whole when making implementation decisions.

#### 2.3.2 Prototyping STEPS 4

Choosing the distributed mesh library with the most suitable abstractions and best performance properties is vital for the success of STEPS 4. This library is the backbone of the whole implementation, providing fundamental data layout and access functionalities, which tightly associates with the criteria discussed above. Besides performance considerations, from a developer’s perspective, the mesh library should also provide a rich and extendable API that can be connected with other STEPS components with ease. Furthermore, while STEPS 4 mainly targets CPUs, the algorithms themselves could in principle be implemented on other hardware architectures and the right abstraction layer should allow a relatively smooth transition towards supporting shared-memory parallelism or GPUs.

To investigate the advantages and drawbacks of different distributed mesh libraries, we used them to implement a series of stand-alone mini-applications to cover a wide range of STEPS functionalities, from simple mesh importing and exporting in a distributed manner, to a functional reaction-diffusion solution integrated with various validations and use case models. These mini-applications were gathered in a library named Zee. Using the Zee library we were able to investigate how different components of STEPS, for example, the operator splitting method, can be implemented on top of different distributed mesh libraries, and to investigate the coding flexibility as well as the performance of our implementations. These investigations provided us essential insight for the choice of a suitable distributed mesh library for STEPS 4, and prototypes for the actual implementation.

We put our evaluation focus on two distributed mesh library candidates, Omega_h (Ibanez and Roberts, 2018) and the DMPlex module from the PETSc library (Abhyankar et al., 2018). Both libraries provide very well-suited features and showed promising performance. The choice for the library, however, also depends on factors beyond pure technical considerations. On the one hand, PETSc seemed a natural choice since STEPS 3’s parallel EField solver already uses PETSc as a backend. Choosing PETSc’s DMPlex would eliminate the need for an extra library, as well as the associated data conversions and transfers between libraries. Additionally, PETSc is an extremely well-known and supported library with a large active community. On the other hand, DMPlex is a minor component in the PETSc framework, supported only by few developers and with a small user community. Since the Zee mini-applications revealed that not all functionalities required in STEPS 4 are currently present in DMPlex, and some of which have considerably low priority on the PETSc development roadmap, our choice had to fall on Omega_h.

Omega_h is a C++14 library providing highly-scalable distributed adaptive meshing primitives. Distributed-memory parallelism is natively supported through MPI, while on-node shared-memory parallelism is supported via Kokkos (Trott et al., 2022), a C++ library that provides abstractions for parallel execution with OpenMP on CPU and CUDA on GPU. Omega_h ensures a fully deterministic execution. Given the same mesh, global numbering and size field, mesh operations produce the exact same results regardless of parallel partitioning and ordering. This does, however, not extend to changing compilers or hardware. Omega_h is being actively developed and is used for a number of ongoing projects. Moreover, its codebase being much smaller than PETSc, it allowed us to have a comprehensive overview of its capabilities. Despite the lack of documentation, the source code is concise and self-explanatory. Contributing to Omega_h has been much easier than it would have been with PETSc. We were for instance able to add support to the MSH multi-part file format version 4 into Omega_h quite easily.

Omega_h’s modern C++ interface was a significant advantage over PETSc as its ease of use allowed us to implement compact yet expressive mini-applications very quickly. We found that the C-oriented API of PETSc makes the library hard to comprehend and is much more error prone than Omega_h’s. Additionally, the data management policies of DMPlex are quite complex and require a deep knowledge of PETSc internals as entity data is not directly exposed to the user as it is in Omega_h. This leads to the code being more cluttered and difficult to maintain.

#### 2.3.3 Solver components and the simulation core loop

Fundamentally, STEPS 4 adopts the same operator splitting solution for reaction-diffusion simulation as in STEPS 3 (Hepburn et al., 2016), but with significant differences in the implementation details due to its distributed nature and other optimization goals.

In STEPS 3, the data and operators are intermixed in the solver, and data that are associated may be stored sparsely due to the data structures inherited from previous STEPS implementations. For instance, the molecule state of a tetrahedron and the states of its neighboring tetrahedrons may be stored far away from each other in memory. This is because the molecule state is stored sparsely in individual tetrahedrons together with other data such as mesh connectivity and kinetic processes. This means operator visits to the molecule state often require significant address jumps across memory, decreasing cache efficiency. The bundle of operators and data also make their optimization cumbersome, as new operator solutions or new data structures can not be implemented directly as independent alternatives.

In STEPS 4 one critical implementation change is the separation and encapsulation of different solver components. The two major components are: SimulationData, the data that represents the current state of the simulation, and the operator collection, which are applied to the data so that the simulation evolves to the next state. The simulation state consists of the molecule state *M*, where the distribution of molecule species is stored and updated, the kinetic process state *K*, which stores and maintains all kinetic processes such as reactions and surface reactions in the simulation and the information of each kinetic process, including the propensity and update dependencies, and finally the voltage state *V* of the mesh if voltage-dependent surface reactions and channels are expressed in the model. The voltage state contains the electrical potential at each vertex of the mesh, as described in Hepburn et al. (2013). The operator collection consists of the operators needed for each step of the simulation core loop, mainly, the reaction SSA operator, the diffusion operator and the PETSc EField operator. As the state data is encapsulated and accessed by operators via a unified interface, new operators can be easily developed and provided to the solver as alternative solutions. The encapsulation of simulation states *M, K* and *V* also allows the state data to be stored contiguously in memory space, thus improving caching efficiency of the solution.

As mentioned in Introduction, while the kinetic processes and their dependency graphs are partitioned and distributed among computing cores in STEPS 3, all mesh elements and their molecule states are duplicated, leading to high memory consumption and communication overhead when dealing with large scale models. In STEPS 4, the mesh itself is partitioned and distributed, thus each computing core only operates on the data for the sub-domain problem of its associated partition. Ghost layers were implemented for partition boundaries so that simulation states of the boundaries can be synchronized through regular data exchange. This solution ensures a relatively consistent memory footprint for any given sub-domain problem with a fixed partition size, regardless of the size of the overall problem.

Figure 1 schematically illustrates the simulation core loop. When the simulation enters the core loop that advances the simulation state from time *T*_*start*_ to *T*_*end*_ = *T*_*start*_ + Δ*T*, the simulation period is divided into multiple time windows, whose period is either determined by a user-defined EField period Δ*T*_*EField*_ if the EField operator is involved, or equals Δ*T* otherwise. We call this the EField time window. Each EField time window is then further subdivided by a period of Δ*T*_*RD*_, where Δ*T*_*RD*_ is determined by the mesh and the diffusion constants of the simulated model. This is the reaction-diffusion (RD) time window.

**Figure 1:**
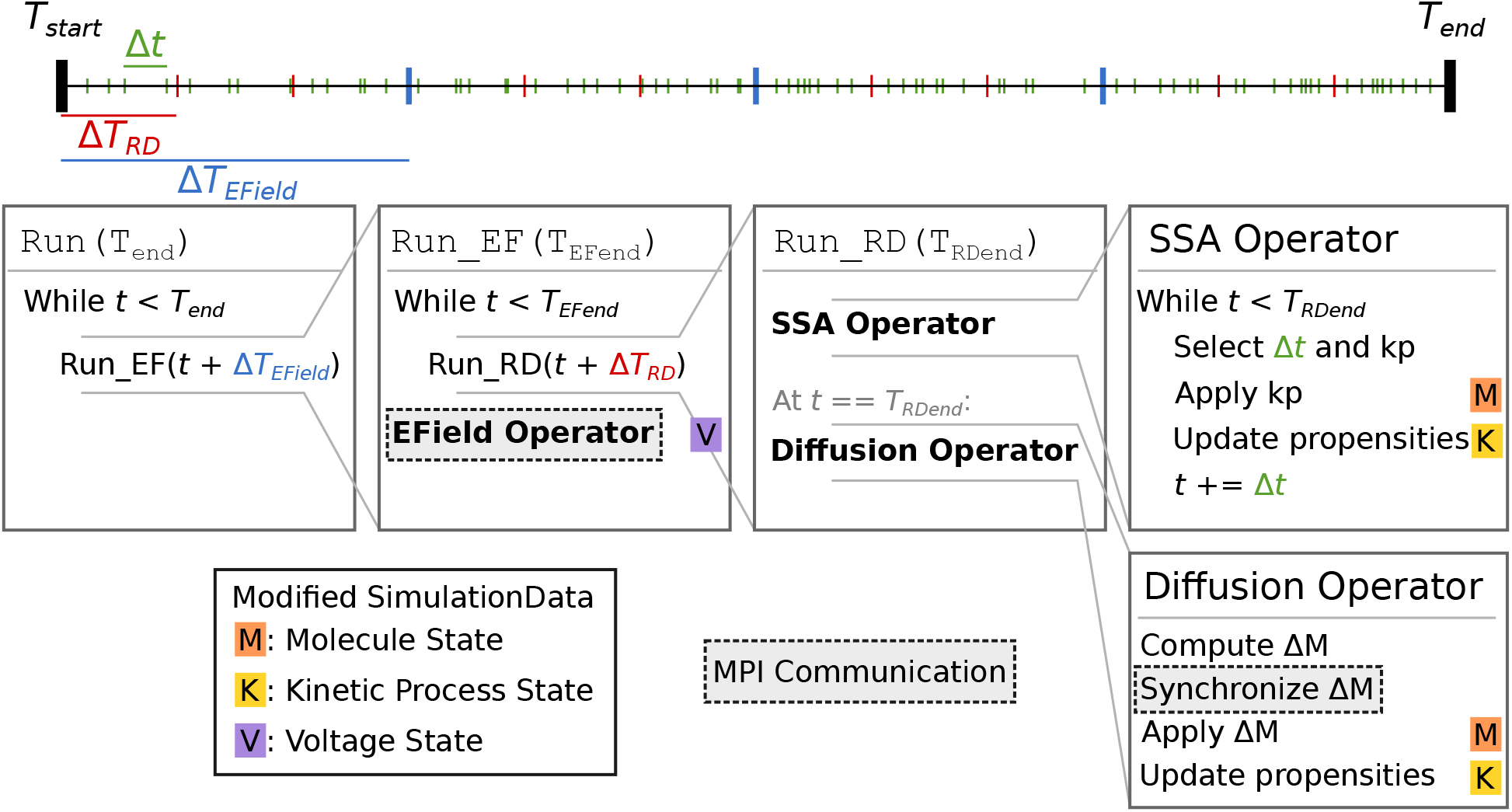
Schematic representation of the STEPS 4 simulation core loop. In this example, when running the simulation from *T*_*start*_ to *T*_*end*_, the simulation time is first split into Δ*T*_*EF ield*_ time windows (blue ticks). Each Δ*T*_*EField*_ time window is further subdivided into Δ*T*_*RD*_ time windows (red ticks). Kinetic process events are represented as green ticks, their number in each time window depends on the propensities of the reactions. Current state time is denoted *t*. The leftmost Run(T_end_) box is the entry point into the core loop, it splits the time in Δ*T*_*EField*_ time windows. The second box (Run_EF(T_EFend_)) runs a full EField time step until *T*_*EFend*_, the end-time that was passed form the first loop. It first subdivides the EField time window in Δ*T*_*RD*_ time windows and calls `Run_RD` for each one. When state time *t* reaches *T*_*EFend*_, it runs the EField operator. The third box (Run_RD(T_RDend_)) represents one RD time window, it is composed of the SSA operator and the diffusion operator. The SSA operator first selects and applies kinetic processes until the state time reaches *T*_*RDend*_; it’s in this loop that the state time is updated. The diffusion operator is then applied: it computes the changes Δ*M* to the molecule state and applies them. Each of these steps can involve the modification of the simulation data. When it does, a letter with a colored background is present to its right. The letter *M* with an orange background signifies that this operation modifies the molecule state; the letter *K* with a yellow background signifies that it modifies the kinetic process state; and the letter *V* on a purple background signifies that it modifies the voltage state. Finally, steps with a darker background and a dashed outline involve MPI communication between processes.

At the beginning of each RD time window, the reaction SSA operator is applied to the simulation data repeatedly. Each time, the SSA operator first randomly selects a kinetic process event *kp* from the kinetic process state *K* and the event time Δ*t* according to the SSA solution described by the operator and the propensities of the kinetic processes. It then applies the molecule changes caused by the event to the molecule state *M*, updates the propensities of all kinetic processes that depend on *kp* in *K*, and advances the simulation state time for Δ*t*. The SSA iteration stops when the state time reaches the end of the RD time window. As explained in (Hepburn et al., 2016) and (Chen and De Schutter, 2017), the SSA operator is executed independently by each MPI rank without the need for any communication.

At the end of the the RD time window, the diffusion operator computes the number of molecules that should diffuse out of each tetrahedron for the time window period Δ*T*_*RD*_. For this calculation the diffusion rates of each diffusive molecule species must be taken into account. The operator then removes them from their original tetrahedrons and redistributes them to their target tetrahedrons. The redistribution is stored in a delta molecule state Δ*M*, which is then synchronized by Omega_h across all simulation ranks. After the synchronization, each rank applies the changes in Δ*M* to *M* for the tetrahedrons it owns, and updates the propensities that are affected by the changes. This completes the operations in a single RD time window.

The solver then repeats this process until the state time reaches the end of the EField time window, at which point the EField operator evolves the voltage state *V* for the period of Δ*T*_*EField*_, based on the electric currents computed from *M* and *K*. This concludes the operations in a EField time window.

If Δ*T*_*EField*_ < Δ*T* the EField time window process is repeated, otherwise the simulation core loop is completed and the user regains the simulation control for data inquiry.

#### 2.3.4 Optimization on kinetic process dependency graph

Efforts have been made to also improve the fundamental solution of the SSA operator, specifically on the optimization of the kinetic process dependency graph. A kinetic process dependency graph describes the update dependency of each kinetic process in the system. Technically, it returns a list of kinetic processes, whose propensities should be updated when a certain kinetic process is selected and applied by the SSA operator. Under the operator splitting framework, the reactions in each tetrahedron are independent until the diffusion operator is applied. Therefore it is possible to subdivide the dependency graph into independent subgraphs and apply the SSA operator to them separately. This independent graph optimization further compresses the targeting domain of the SSA operator, providing potentially substantial gains in simulation performance.

An example of the optimization for a small model is depicted on Figure 2. This model has two tetrahedrons, each with three volume reactions. One tetrahedron also contains four surface reactions. Each colored node in the figure represents a kinetic process. An arrows goes from one node to the other if the occurrence of an event of the first entails a change in propensity of the second. The whole dependency graph of the model can therefore be subdivided into two independent subgraphs, in red and blue as shown in the figure. Each subgraph can be evolved freely by a SSA operator without the other’s interference in a RD time window period. Note that the effect of this optimization depends on the duration of the RD time window period, as well as the rates of the reactions. Generally speaking, this optimization favors simulations with high molecule concentrations and highly active reactions, but disfavors simulations with low molecule concentrations and less active reactions.

**Figure 2:**
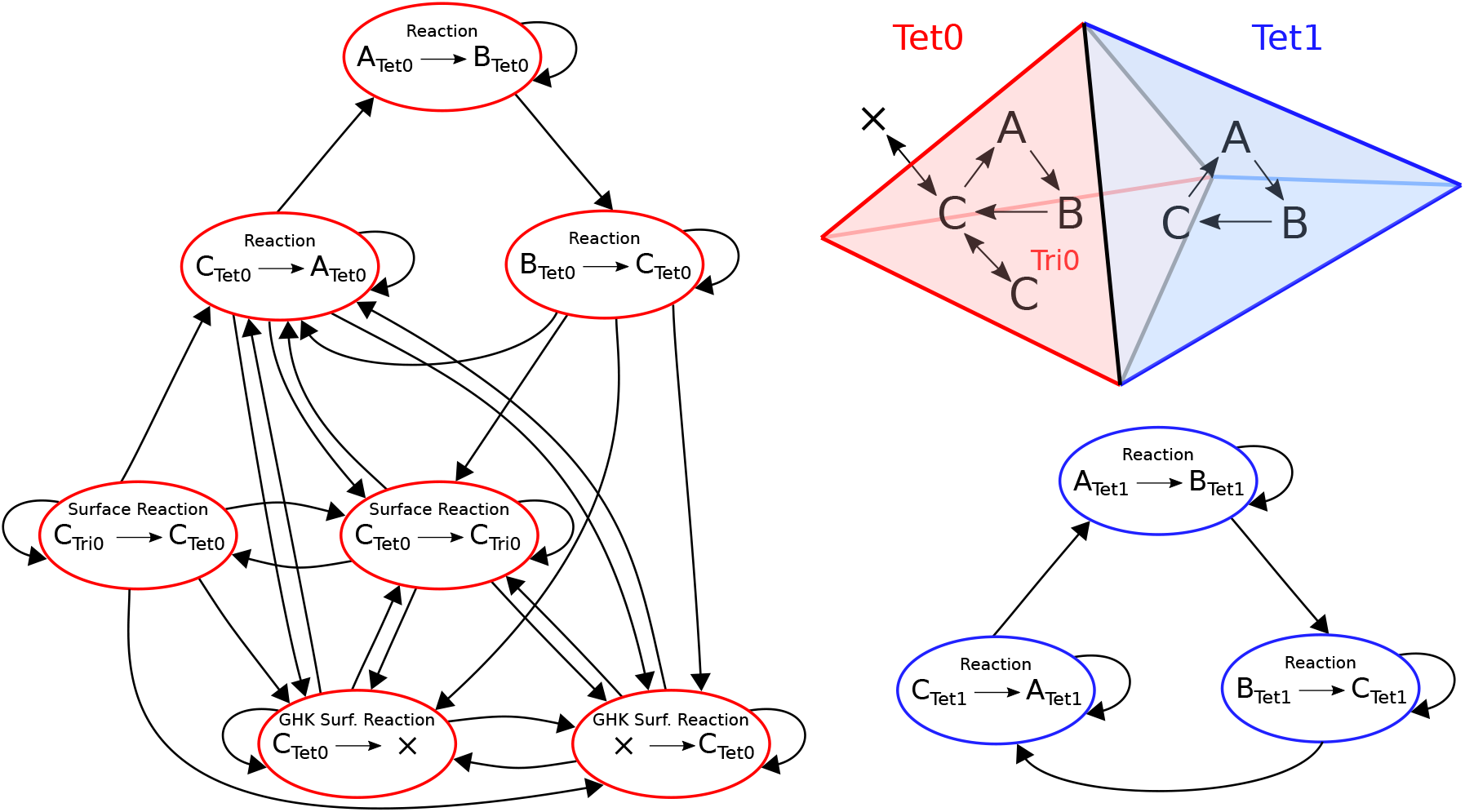
Structure of the reaction dependencies graph on a mesh with two connected tetrahedron labeled 0 and 1. The top right part of the figure represents the two tetrahedron and the reactions they contain. Both tetrahedrons contain reactions *A* → *B, B* → *C* and *C* → *A*. Tetrahedron 0 also contains four surface reactions: species C can be transferred back and forth to a triangle (*C*_*tet*_ → *C*_*tri*_ and *C*_*tri*_ → *C*_*tet*_); and species C can cross the membrane back and forth as a GHK current (since the amount of *C* outside of tetrahedron 0 is not modeled, it is equivalent to creating and removing species *C*). In the main figure, each colored node represents a kinetic process. An arrows goes from one node to the other if the occurrence of an event of the first entails a change in propensity of the second. In blue and red are the extracted connected components of the graph.

#### 2.3.5 EField solver improvements

Generally, in order to obtain the most accurate results and the best performance, solver and preconditioner need to be tailored to the particular simulation. Previously STEPS 3 used by default the Conjugate Gradient iterative solver (CG) and the Geometric Algebraic Multigrid (AMG) preconditioner. However, performance tests have consistently shown that they do not scale well for large problems. Thus, for STEPS 4 we replaced solver and preconditioner with the widely used Pipelined Conjugate Gradient method (KSPPIPECG) and the Point Block Jacobi preconditioner (PCPBJACOBI), respectively. The same configuration was also applied to STEPS 3 as the new default option. We have not performed a thorough investigation on solvers and preconditioners as it was out of the scope of the present manuscript.

Another improvement is the distribution of PETSc vectors and matrices for the EField computation. STEPS 3 distributes them equally among computing cores without considering if the mesh elements represented by the matrix partition are owned by the same core. This causes owner mismatches between the EField solution data and the reaction-diffusion solution data, which need to be resolved by expensive cross process data exchanges. In order to avoid this issue, STEPS 4 assembles the vectors and matrices so that each processor only takes care of the degrees of freedom corresponding to the sub-part of the mesh that is owned locally on this processor. This greatly increases data locality and performances since reaction-diffusion and the EField solvers exchange data only locally.

#### 2.3.6 Mesh generation, partition and annotation

Due to the distributed nature of STEPS 4, a new mesh importing, partitioning and annotation solution has been added, supported by the Omega_h and Gmsh (Geuzaine and Remacle, 2009) libraries. Currently it supports importing Gmsh .msh meshes, with or without pre-partitioning. If the mesh is not pre-partitioned, the Omega_h backend in STEPS 4 performs a recursive bisection partitioning on the mesh at MPI rank 0, then distributes each mesh partition to the owner rank. If the mesh is pre-partitioned in Gmsh, each rank imports its associated partition directly from the file and then establishes the connectivity to other processes. In both cases, the number of MPI processes used in a STEPS 4 simulation should always be a power of 2. It is also worth mentioning that the two solutions apply different partitioning schemes, which may affect simulation performance drastically depending on the mesh morphology.

Another significant change on mesh preparation is the annotation of physical entities of the mesh such as “compartments” representing volumetric regions such as the cytosol and extracellular space, and “patches” representing surface regions such as the cell membrane and intracellular membranes. In STEPS 3, the user provides a list of tetrahedron/triangle indices to create a compartment/patch of the mesh. The index lists are either generated on the fly during the simulation, or generated in advance, stored externally then provided to the simulation. The user is expected to take full responsibility of the generation and maintenance of these lists. In STEPS 4, the annotation of compartment/patch is embedded into the mesh file itself by utilizing the physical tag functionality in Gmsh. As a general procedure, the user can create physical entities on the mesh using the Gmsh application and provide a string as the physical tag for each of the entities. These string tags can then be used as identifiers to create the compartments and patches in STEPS 4. If the anticipated boundaries of the compartments/patches are irregular, but can be described by a combination of enclosed polyhedral surfaces, then the user can also utilize the gmsh backend implementation of the PolyhedronROI^1^ utility to embed the physical tags into associated mesh elements.

#### 2.3.7 Coupling with other STEPS components

Setting up a simulation in STEPS 4 is mostly done in the same way as in STEPS 3: it involves the declaration of a biochemical model and a description of the geometry in which the model will be simulated. The biochemical model is composed of species, channels, reactions, diffusion rules and currents that are grouped by volume or surface systems. Although most of the biochemical modeling features available in STEPS 3 are also available in STEPS 4, surface diffusion rules are not yet supported. Internally, the same classes are used for declaring a biochemical model in STEPS 3 and in STEPS 4. While in STEPS 3, tetrahedral meshes were managed with the `TetMesh` class, a different class (`DistMesh`) was added for distributed meshes in STEPS 4. This class inherits from the same `Geom` base class as `TetMesh` but acts as a wrapper around the `Omega_h::Mesh` distributed mesh class. Classes related to the declaration of compartments (`DistComp`), patches (`DistPatch`) and membranes (`DistMemb`) in a distributed mesh are also different from the ones used in STEPS 3. Most notably, as explained in the previous section, while tetrahedral compartments in STEPS 3 are usually built from a list of tetrahedron identifiers, STEPS 4 makes use of physical tags in distributed meshes to create distributed compartment and distributed patches. On solver creation, the `DistTetOpSplit` distributed solver class in STEPS 4 initializes the relevant data structures from the biochemical model and geometry description classes. Although this type of initialization through the python API corresponds to the most frequent use case, the distributed solver can also be used and initialized directly in C++, without requiring the creation of a STEPS biochemical model and geometry classes.

#### 2.3.8 Python API changes due to the distributed nature

As described above, the main difference between a serial or parallel model and a distributed model consists in the declaration of compartments and patches. In order to use the `DistTetOpSplit` solver in STEPS 4, one must first create an instance of the `DistMesh` class. Compartments and patches are then usually created by using physical tags in the mesh: the tetrahedrons/triangles tagged with the name of the compartment/patch will be used to build it. The following example shows the creation of a compartment and a patch in a distributed mesh:

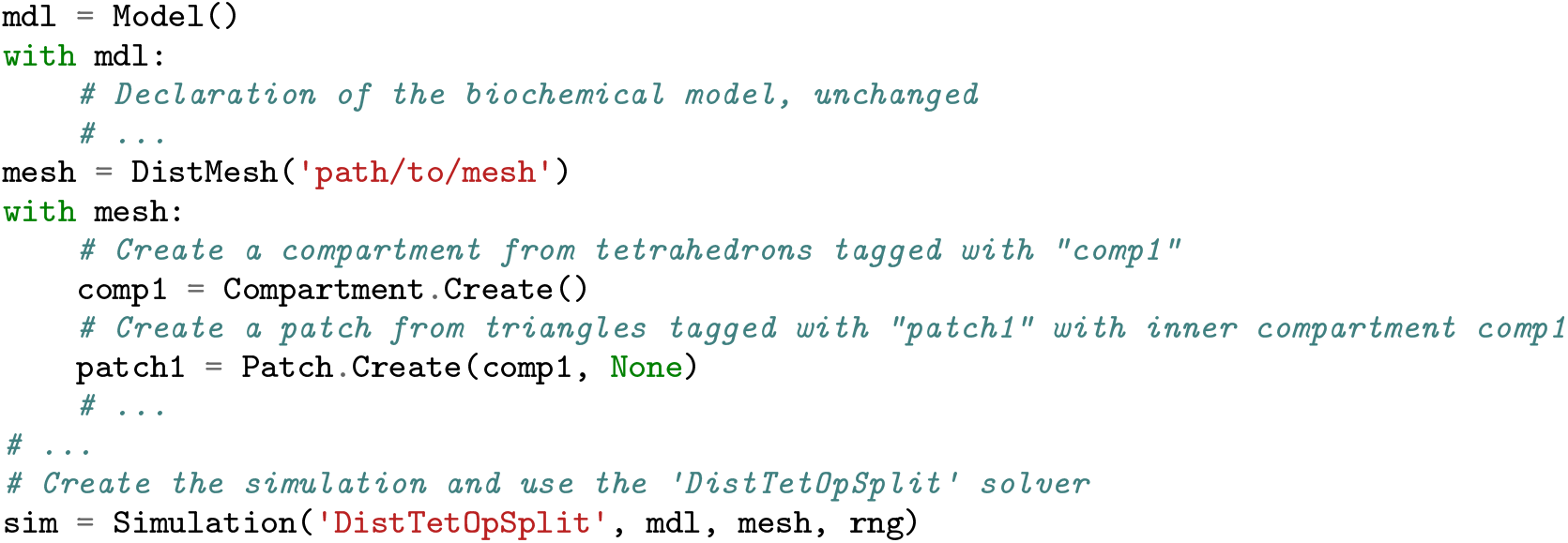

Note that it is also possible to use lists of tetrahedrons to create a compartment. In that case, the compartments and patches creation code is identical to STEPS 3. Finally, automatic data saving code is unchanged. Detailed documentation is available^2^.

### 2.4 Validation strategy

In order to ensure accurate results, STEPS 4 is validated on a series of published models. The faster validations are integrated into the STEPS release and used in continuous integration while the others are available in the STEPS validation repository ^3^. However, since these models are stochastic models that are designed to run in a reasonable amount of time, they each contain a small tolerance that could mask minor numerical inaccuracies. So as to rigorously test our new methods and implementations in STEPS 4 and ensure even no small loss of numerical accuracy, we go further in this manuscript and investigate STEPS 4 in a series of models, comparing either to STEPS 3 results or analytical solutions to a high degree of accuracy.

Validating stochastic simulation solutions has always been a difficult challenge. Often, analytical solutions exist only for a few trivial problems and, even in those cases, the stochastic nature of the simulator may make results fluctuate around the analytical solution depending on the particular seed provided to the random number generator (RNG). Unfortunately, fixing the seeds and numerically comparing STEPS 3 and 4 results is not a meaningful strategy since the two simulators use RNG streams in different ways. Thus, we validate STEPS 4 in a statistical sense.

#### 2.4.1 Statistical analysis

We extract meaningful statistical data from multiple realizations with different RNG seeds and compare either with STEPS 3 results or the analytical solution when available.

The general steps are:

- Record relevant trace results such as the voltage traces in a particular location in the mesh from multiple realizations of STEPS 3 and 4 simulations.
- Refine traces to extract key features of the simulation. E.g. the frequency of a spike train.
- Collect refined features among the various simulation runs and statistically compare STEPS 3 and 4.

The choice of what must be recorded and what are the relevant features depends on the particular model at hand.

We choose the well-known two independent samples Kolmogorov-Smirnov test (KS test) for our statistical comparisons between STEPS 3 and STEPS 4, utilizing the Scientific Python (SciPy) library. The null hypothesis in our KS tests is that the two samples come from the same distribution. Perhaps a common misconception of this test is that a p-value below 0.05 means that the null hypothesis must be rejected and, therefore, the distributions are different. In fact, when comparing two identical distributions the p-value is expected to be uniformly distributed on [0,1], and so if this test is repeated many times one would expect to see a p-value below 0.05 5% of the time. In our tests, where multiple distributions are compared within one model, we reject the null hypothesis only if there is strong evidence that p-values are consistently low, evidenced by significantly more than 5% of the p-values generated being below the 0.05 level.

Conversely, when traces are relatively smooth and the features are few, we study directly the confidence intervals at the customary 95% confidence level. In this case, we reject the null hypothesis if the mean of the STEPS 3 traces does not lie in the confidence interval of the STEPS 4 traces or vice versa.

## 3 Results

### 3.1 Validations

As STEPS 4 contains multiple operator components targeting different sub-systems, such as molecular reaction-diffusion and EField, we carefully select the models and independently validate each component before testing the whole implementation on a complex, real case scenario.

#### 3.1.1 Validations of the reaction-diffusion solver

We extend the validation pack described in Hepburn et al. (2012) and Hepburn et al. (2016) to validate the reaction-diffusion solver and the basic functionalities of other data structures introduced in the new code implementation. STEPS 4 passes all of them. For the sake of brevity and since the reaction-diffusion operator splitting solution remains theoretically the same as in STEPS 3, we hereby skip the detailed analysis of these validations.

#### 3.1.2 Validations of the EField solver

To validate the EField solver we use the Rallpack models described in Bhalla et al. (1992).

- Rallpack 1 simulates a simple uniform unbranched passive cable. Since no randomness is involved in this validation, STEPS 4 results are compared directly to the analytic solution.
- Rallpack 2 model is superfluous and can be skipped here as it is the same model as Rallpack 1 but with a branching morphology. This mathematical morphological description is also in practice very difficult to capture realistically in a mesh.
- Rallpack 3 examines the interaction between the EField system and the stochastic channel activities of the well-known Hodgkin-Huxley model (Hodgkin and Huxley, 1952). No analytical solution is available for this test, thus we compare STEPS 3 and 4 solutions using the statistical validation framework illustrated in Statistical analysis.

##### Rallpack 1

Rallpack 1 (Bhalla et al., 1992) focuses on the validation of the EField solver and is one of the most basic examples one can think of. It consists of a leaking, sealed straight cable with a current injection (*J*) at *z*_*min*_. For the sake of brevity, Rallpack 1 setup is depicted in Figure 3 **(A)**. Table 1 provides the parameters.

**Table 1:**
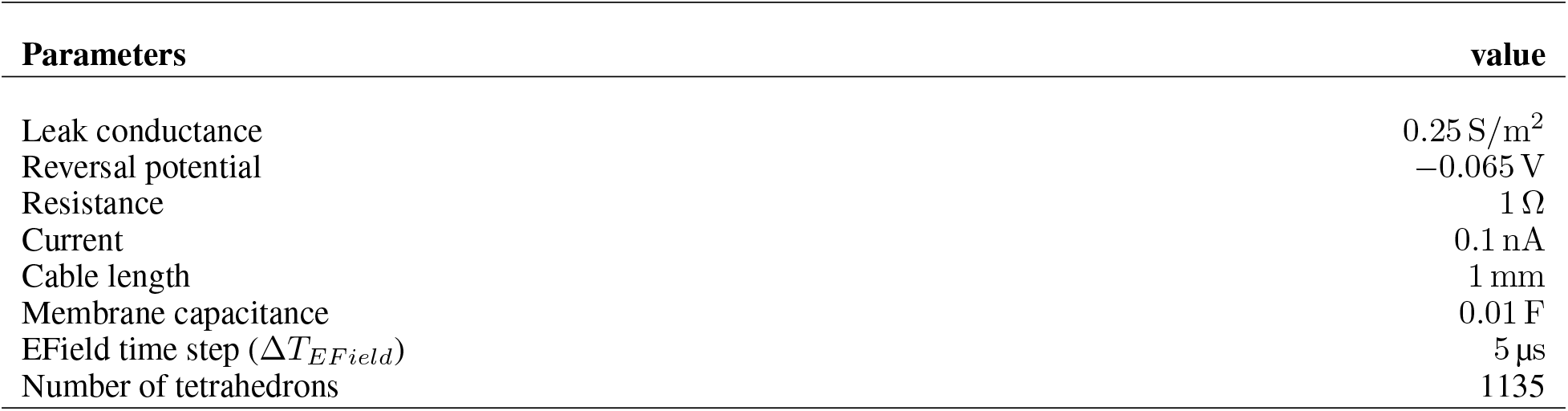
Parameters for Rallpack 1.

| Parameters | value |
| --- | --- |
| Leak conductance | 0.25 S/m<sup>2</sup> |
| Reversal potential | -0.065 V |
| Resistance | 1 $\Omega$ |
| Current | 0.1 nA |
| Cable length | 1 mm |
| Membrane capacitance | 0.01 F |
| EField time step ($\Delta T_{EField}$) | 5 μs |
| Number of tetrahedrons | 1135 |

**Figure 3:**
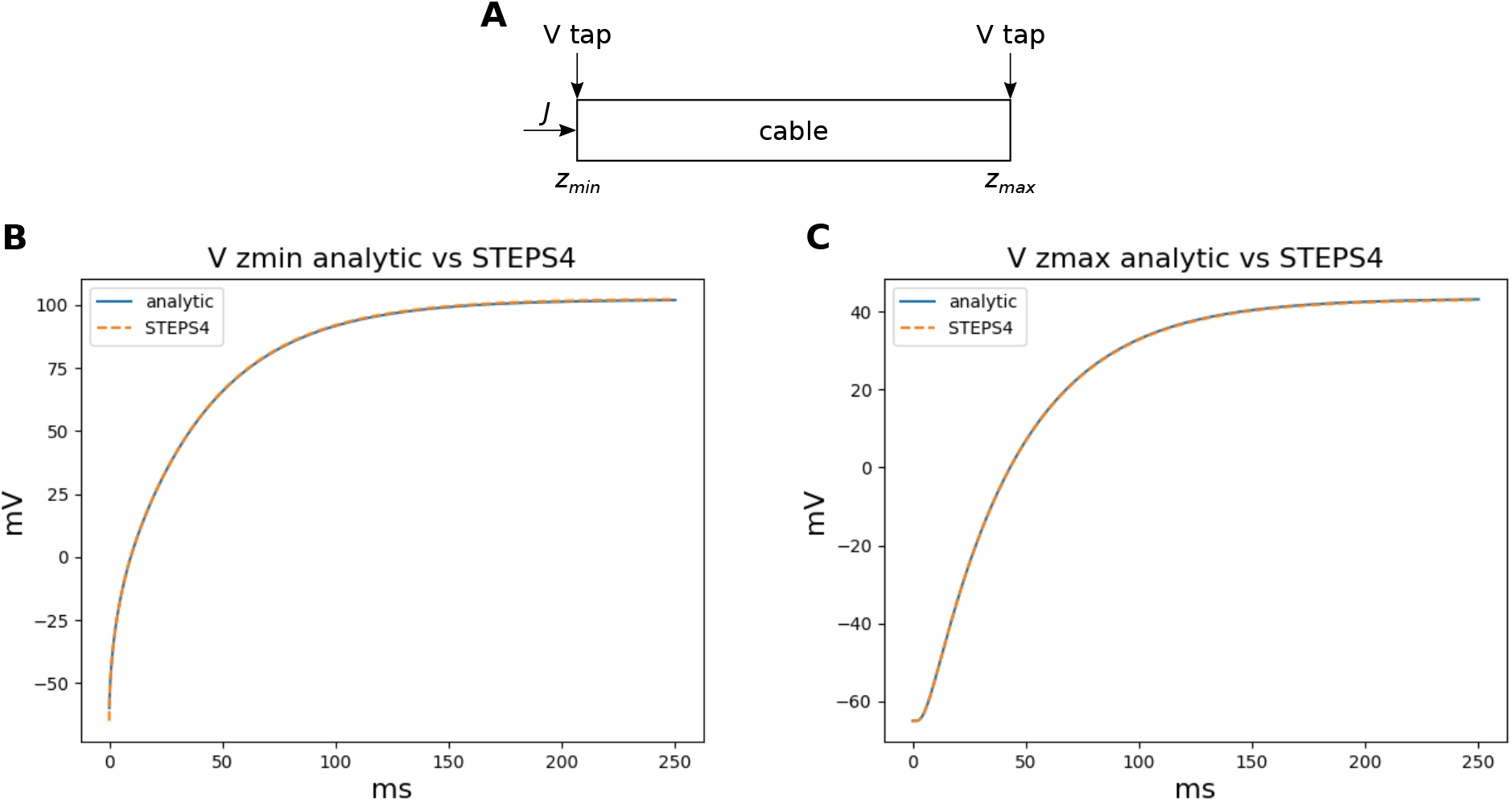
**(A)** illustrates the general setup for Rallpack 1. **(B)** and **(C)** compare STEPS 4 voltage traces with the analytical solution at the extremes of the cable respectively. Results overlap with mse: 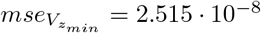 and 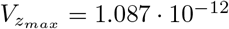.

Without loss of generality, we can focus on the voltage traces at the extremes of the cable because the equations are linear and all the intermediate solutions are super-positions of the results at the extremities.

Since this simple problem presents an analytical solution and no source of randomness, it is the perfect candidate to benchmark the STEPS 4 EField solver passive properties. A leak channel is introduced on every surface triangle. This is slightly different from the analytical solution setup where the leak is uniformly distributed along the cable. However, the effects should be negligible if the mesh is sufficiently refined.

Figure 3 visually compares STEPS 4 results with the analytic solution. As expected, the curves overlap and the mean square errors (mse)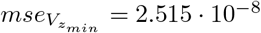 and 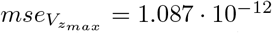 are quite small compared to the traces. It is interesting to note that STEPS 3 presents almost exactly the same results. In fact, when comparing STEPS 3 with STEPS 4 on the same mesh, the mse is 1*e* − 18 for both 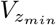 and 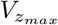 and is due to numerical precision.

Convergence through mesh refinement proceeds as expected with an initial steep drop followed by a plateau at numerical precision. For the sake of brevity, it is left in the Supplementary Material Section S1.1.1.

##### Rallpack 3

Rallpack 3 is an active model that builds on Rallpack 1 by adding Hodgkin-Huxley sodium and potassium channels, and is simulated on the same simple, uniform, unbranched cable geometry. The model tests ion channel activation as well as spike propagation. As for Rallpack 1, voltage is recorded and analyzed at the two ends of the cable and referred to as 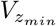 and 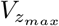. Differently from Rallpack 1, Rallpack 3 presents sources of randomness and the problem cannot be solved analytically. A statistical analysis is employed to study this simulation and validate the code.

The two sample sets consist each of 1000 simulation runs performed with STEPS 3 and 4 respectively. As for Rallpack 1, we record voltages at the extremes of the cable 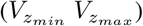 (the raw traces). Figure 4 illustrates two realizations of the model.

**Figure 4:**
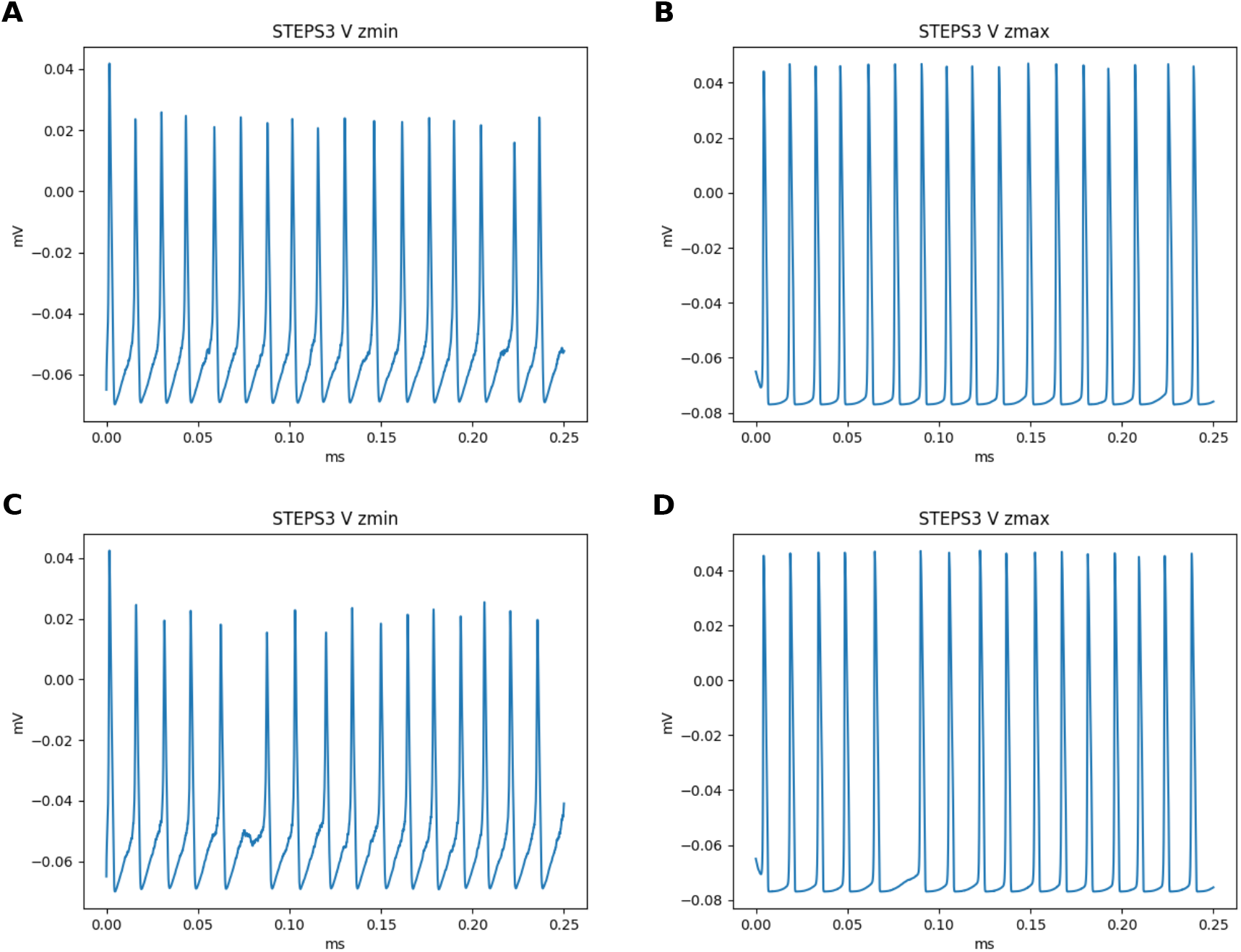
The panels present the traces at *z*_*min*_ and *z*_*max*_ for two different realizations of the simulation with STEPS 3. Panels **(A)** and **(B)** illustrate a run where peaks are present. Conversely, the panels **(C)** and **(D)** present a run with a failed spike and thus a skipped peak. The simulated time span is 0.25 s. At *z*_*min*_ a higher peak (∼0.04 V) is followed by a constant spike train while at *z*_*max*_ spikes result all similar to each other and have the same height as the first, bigger peak at *z*_*min*_. Valleys are at ∼-0.065 V and the spike frequency is ∼69 Hz. Since Rallpack 3 is run on a simple conducting cable, the frequency is the same at both ends.

The voltage trace at *z*_*min*_ presents a higher peak at ∼0.04 V followed by a constant spike train which just surpasses V. Conversely, the spike train at *z*_*max*_ has no bigger spike at the beginning (in fact the opposite is true: the first spike is slightly smaller than the others) and their values is above 0.04 V. For both traces valleys are at ∼-0.065 V and frequencies are ∼69 Hz. Over the total simulation time of 0.25 s, usually, 17 peaks appear. However, sometimes a peak or two can be skipped due to failed spikes as illustrated by the second row of results in Figure 4. It is interesting to note that this behavior can occur only with a stochastic simulator.

Given that traces are spike trains with, possibly, a single greater initial peak, the key features extracted and statistically analyzed are:

- peak height;
- peak timestamps;
- the most relevant frequency peak, extracted using the Fast Fourier Transform (FFT);
- number of peaks.

The null hypothesis is that STEPS 3 and 4 simulation results come from the same population, in other words, the simulations are identical. We use the KS test to confute it with the customary 95% confidence level. Figure 5 depicts the p-values and their distributions for timestamp and height for the 17 peaks on 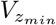 and 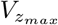 Overall, the p-values look to be distributed uniformly on the interval [0, 1] for all the graphs. This is what we expect to see if comparing two identical distributions, as described in Statistical analysis. Therefore, we cannot refute the null hypothesis, and we accept that the two samples are taken from the same population.

**Figure 5:**
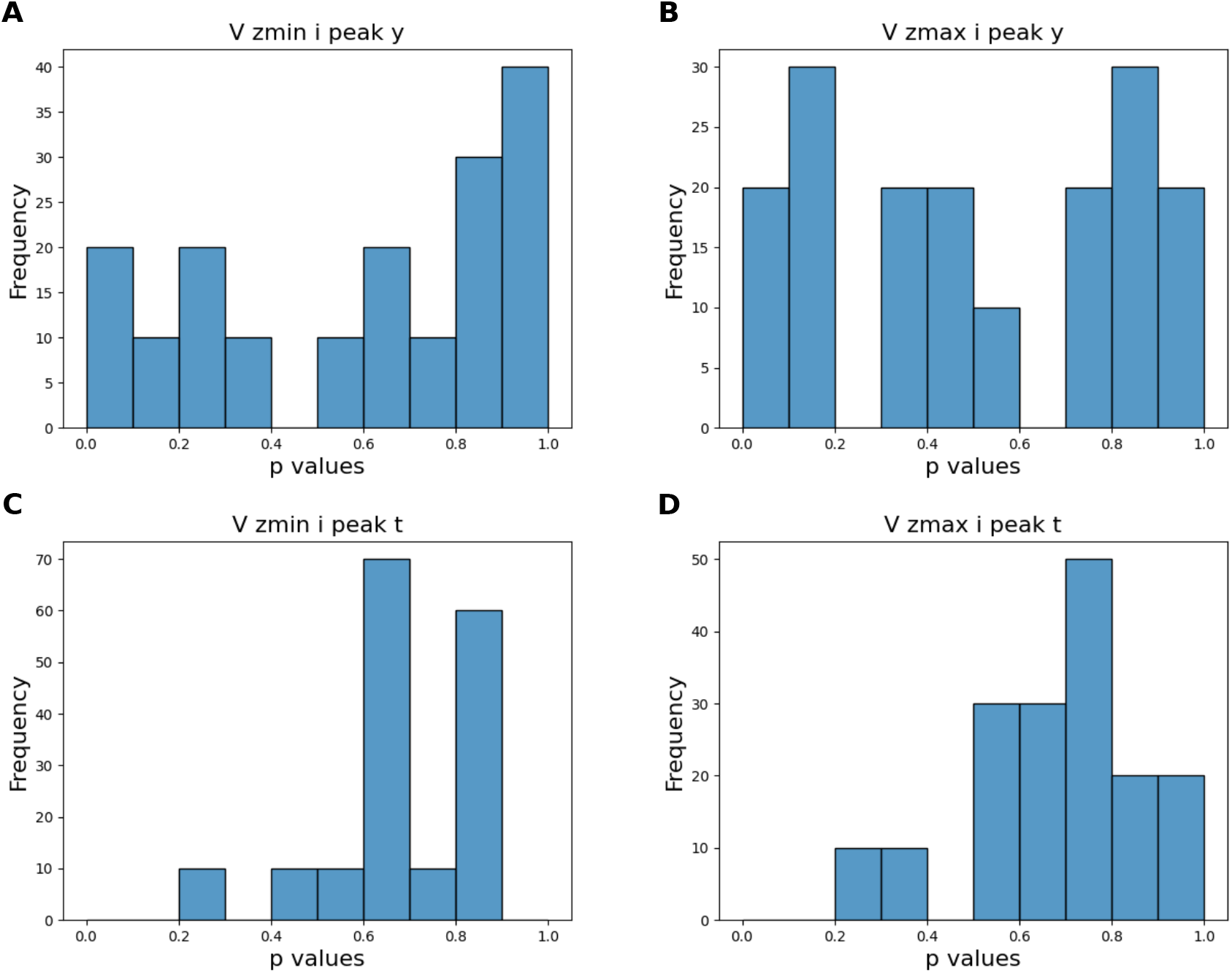
Distribution of the 17 p-values for peak height and timestamp at 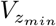 and 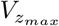. They all present a uniform distribution. A few peaks do not pass the usual p-value test at 0.05. That is expected since, by definition, two samples taken from the same population, on average, do not pass the test once every 20 tests. Since their distribution is overall uniform we refute the null hypothesis and accept that the two samples are taken from the same population.

#### 3.1.3 Full validation

Finally, we validate the full STEPS 4 simulator with a complex scenario, which combines reaction-diffusion and EField features and their possible interactions.

##### The calcium burst model

The previously published calcium burst model (Anwar et al., 2013) is selected for the full validation. It contains most of the modeling features supported by STEPS 4, such as regular molecule reaction-diffusion events, ligand-based channel activation and electric potential dynamics. Thus, it is the perfect candidate to validate STEPS 4 as a whole. Minor modifications are applied to the original model in Anwar et al. (2013) in order to run on a full dendritic mesh, as opposed to the sub-branch mesh used in previous studies. For the sake of clarity, Figure 6 illustrates its dendritic morphology. The full dendritic mesh was created from reconstruction retrieved from NeuroMorpho.Org (Ascoli et al., 2007), data ID: NMO_35058 (Anwar et al., 2014)^4^. The calcium burst model is also used to analyze the performance of the implementation in the Performance section.

**Figure 6:**
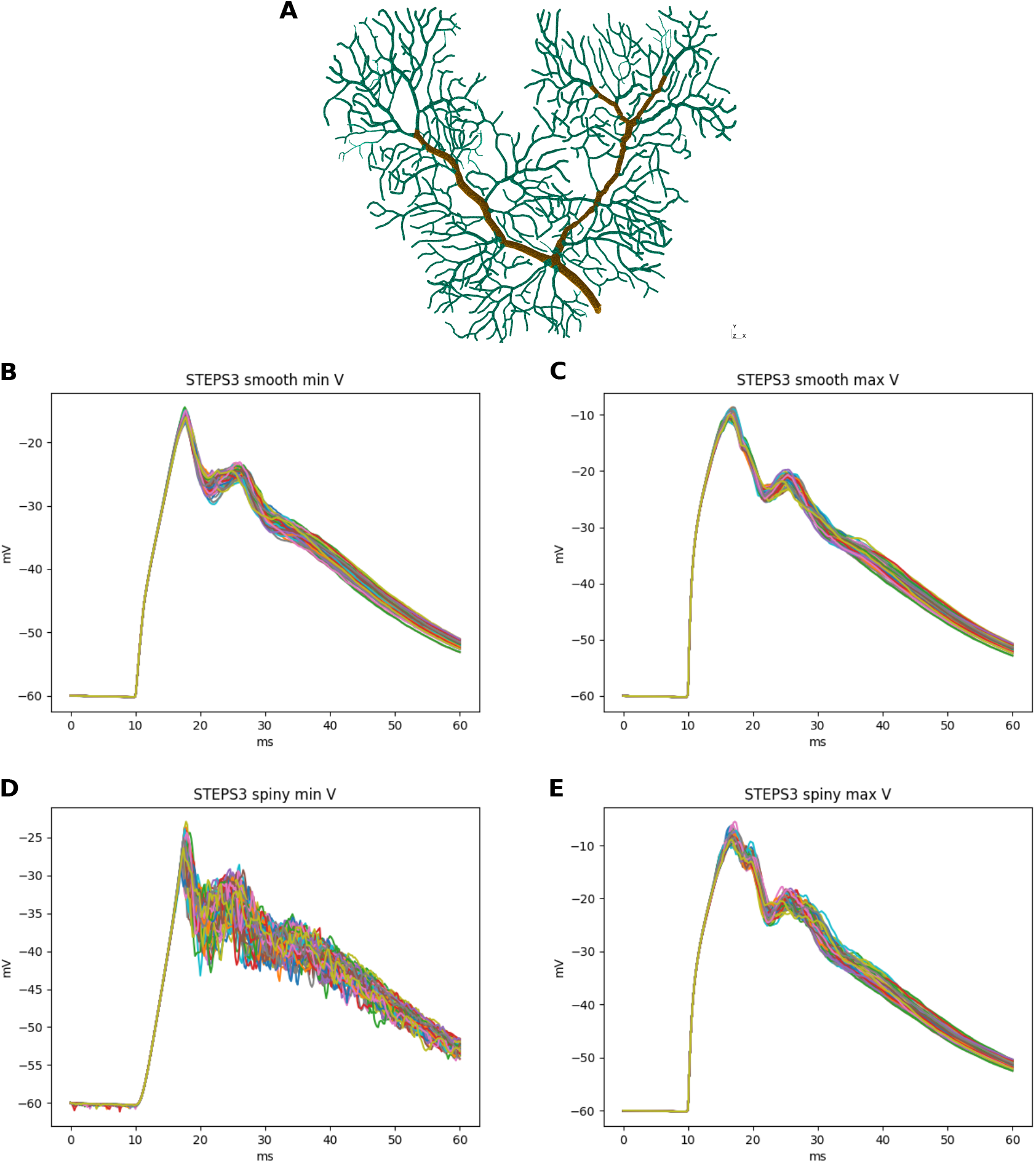
**(A)** illustrates the Purkinje dendrite mesh reconstruction for both the Calcium burst background and complete models. The mesh consists of 853,193 tetrahedrons. Dendrite elements are further classified and annotated into two components, representing the smooth dendrite and the spiny dendrite of a Purkinje cell. Different channel density configuration is assigned to each component in the complete model. The remaining **(B), (C), (D)**, and **(E)** are raw traces of minimum and maximum voltages on smooth and spiny surfaces for 100 runs of STEPS 3. After the first depolarization at ∼18 ms the systems starts to behave stochastically. It also presents another, smaller peak at ∼28 ms before slow repolarizing. The spiny minimum is particularly noisy.

The raw traces used for this analysis are minimum and maximum voltages on smooth and spiny membranes. Figure 6 presents these traces for 100 runs of STEPS 3. After the first depolarization burst at ∼0.018 ms another, smaller peak appears at ∼0.028 ms before slowly repolarizing.

As for Section 3.1.2, our null hypothesis is that the two simulators run the same simulation and results are picked from the same population. Given the low number of peaks and the relatively small stochastic effects, we try to confute this statement computing the confidence intervals of the averages of the traces at the typical 95% probability. By definition, the confidence intervals mark a region where the trace average lies with 95% probability. Thus, if the average of the traces of the STEPS 3 set does not lie in between the confidence intervals of the STEPS 4 set or vice versa we reject the hypothesis. Figure 7 presents averages and confidence intervals of the minimum and maximum of the voltages on smooth and spiny membranes. Confidence intervals are extremely narrow due to the regularity of the traces. Since STEPS 4 average lies in the confidence interval of the STEPS 3 simulation set and vice versa we cannot reject the null hypothesis and we consider STEPS 4 validated even in this complex scenario.

**Figure 7:**
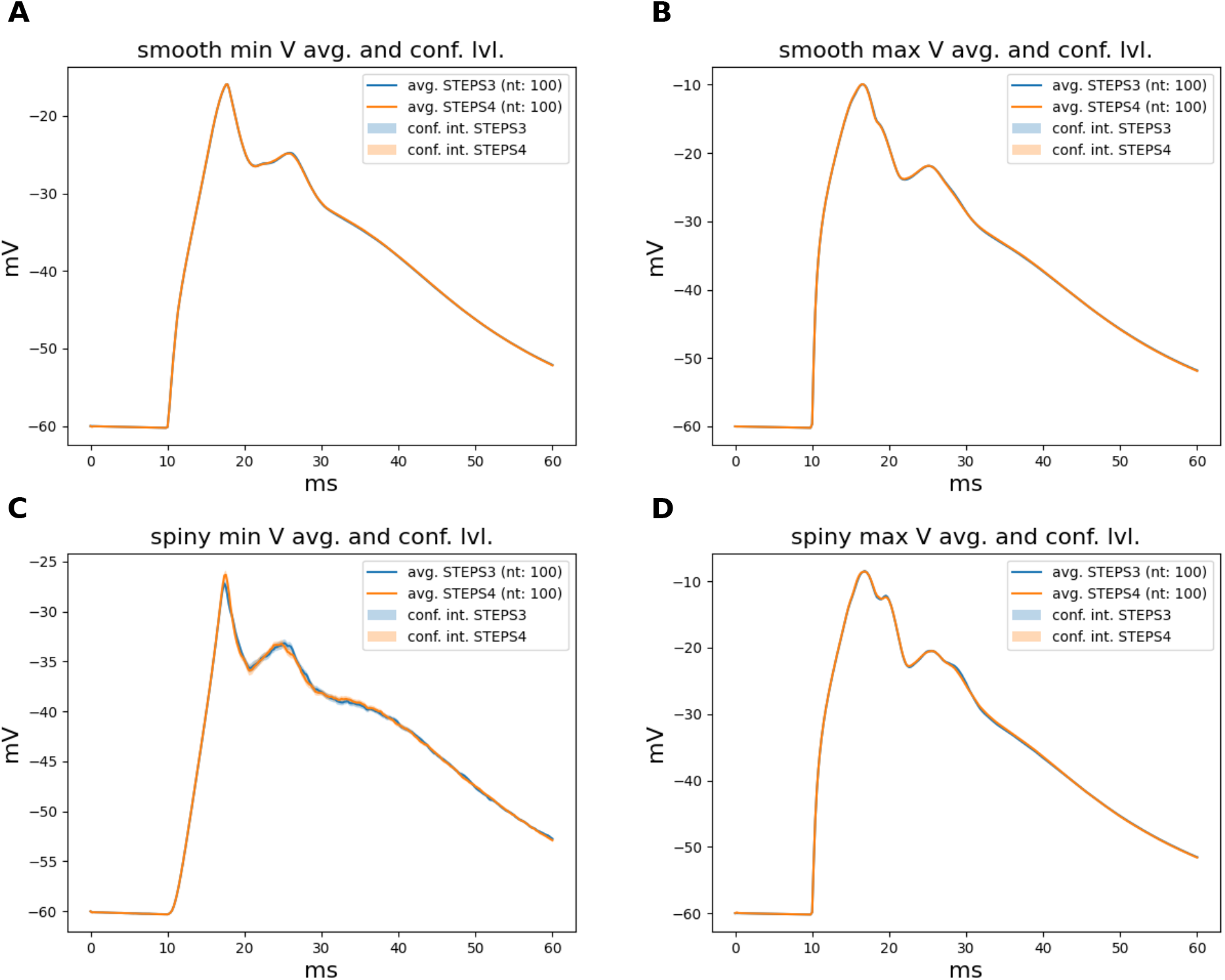
The subplots present average and confidence intervals of STEPS 3 and 4 simulation sets. The traces averages overlap almost perfectly and the confidence intervals at 95% probability are extremely narrow due to the regularity of these results. Since the average of one set lies in the confidence intervals of the other and vice versa we cannot confute the null hypothesis and we consider STEPS 4 validated even in this complex scenario. *nt* stands for Number of Tests.

### 3.2 Performance

We evaluated the performance of the implementation using three models with gradually increased complexity to cover the use cases from a wide range of research interests. The first one is a simple reaction-diffusion model on a simple cuboid mesh. In the second model, we simulate the background activities of the calcium burst model to investigate the performance of the reaction-diffusion solution on complex Purkinje cell morphology. Finally, in the third model we simulate the complete calcium model to study how the combined solution performs with a real world model. The simple model and the calcium burst background model have previously been used to study performance and scalability of the reaction-diffusion operator splitting solution in STEPS 3 (Chen and De Schutter, 2017). As the implementation has been improved since the initial implementation, and the hardware used in the previous research is now outdated, new simulation series of these two models are performed to acquire up-to-date results for comparison. The parallel performance of the combined solution with the complete calcium burst model has not been reported previously. The independent graph optimization is disabled for these simulation series to provide fair comparison between models.

#### 3.2.1 Benchmarking Setup

All simulation benchmarks were run on the Blue Brain V (BB5) supercomputer hosted at the Swiss National Computing Center (CSCS) in Lugano, Switzerland. A complete description of the hardware and software configuration details of the BB5 system are provided in Table 2. All benchmarks were executed in pure MPI mode by pinning one MPI rank per core. As the number of cores used for simulation needs to be a power of 2 (see Section 2.3.6), for each series of benchmark we first choose an initial core count as a baseline and then double the core count.

**Table 2:** Hardware and Software Configurations of the Blue Brain 5 (Phase 2). Supercomputer

|  |  |  |
| --- | --- | --- |
| Hardware | System | HPE SGI 8600 |
|  | Compute Node (880×) | 2× Intel Xeon Gold 6248 Cascadelake @2.5GHz (20 physical cores per CPU) |
|  | Memory | 384GB of main memory (12 × 32GB DDR4-2933 DIMMS) |
|  | Network | InfiniBand EDR 100Gbps / Fat-tree topology |
|  | Accelerator | GPFS/ IBM Spectrum Scale Filesystem (6.2PB) |
| Software | Compiler | GCC C++ compiler 9.3.0 |
|  | Operating System | Red Hat Enterprise Linux Server 7.9 |
|  | MPI | HPE MPI (SGI MPT) 2.25 |
|  | Python | 3.8.3 |
|  | Linked Libraries | PETSc 3.14.1, Omega_h 9.34.6, Intel MKL 2018.3, Eigen 3.3.8, SUNDIALS 2.7.0, mpi4py 3.1.3, NumPy 1.21.4, GMSH 4.9.0 |

The code instrumentation for the performance measurement in STEPS is performed through an *Instrumentor* interface. This is a light wrapper that allows for marking/profiling code regions of interests either by calling a start/stop method or by *C++* Resource Acquisition Is Initialization (RAII) style. Various backends are used by this interface, in particular in this work we use Caliper 2.6 (Boehme et al., 2016), and LIKWID 5.2.0 (Treibig et al., 2010).

For each benchmark configuration, we repeat the simulation 30 times, and show the average results in the figures. The standard deviations of the results are reported as the error bars for each data point in the figures. Per-core memory consumption of each simulation is also measured using the *psutil* Python module (Rodola, 2020) and reported. The comparisons are mainly conducted between STEPS 3 and STEPS 4. For the scalability studies, we also compare the results with the theoretical ideal speedup scenarios. We further investigate the contribution and scaling properties of operator components in STEPS 4, namely, the SSA operator, the diffusion operator and the EField operator, by measuring their individual speedup as well as the proportion in the overall simulation time cost.

#### 3.2.2 The Simple Model

We reuse the simple model in Chen and De Schutter (2017) which consists of 10 diffusing species with different initial molecule counts within simple cuboid geometry with 13,009 tetrahedrons. These species interact with each other through 4 different reversible reactions with different rate constants. The details of the model can be found in Table 3. We choose 2 cores as the performance baseline and increase the core count to 2^11^ = 2048 as the maximum. Note each core has less than 10 tetrahedrons with this maximum, at which point it is unlikely that the simulations remain scalable. However, the result is still interesting as it illustrates the behavior of our solution under extreme scaling scenarios.

**Table 3:** Species and reactions as well as the initial configuration of the simple model.

| Species | Diffusion Coefficient ($\mu\text{m}^2/\text{s}$) | Initial count |
| --- | --- | --- |
| A | 100 | 1,000 |
| B | 90 | 2,000 |
| C | 80 | 3,000 |
| D | 70 | 4,000 |
| E | 60 | 5,000 |
| F | 50 | 6,000 |
| G | 40 | 7,000 |
| H | 30 | 8,000 |
| I | 20 | 9,000 |
| J | 10 | 10,000 |
| Reaction | Rate Constant |  |
| $A + B \rightleftharpoons C$ | $k_f : 1,000(\mu\text{M} \cdot \text{s})^{-1}, k_b : 100\text{s}^{-1}$ | |
| $C + D \rightleftharpoons E$ | $k_f : 100(\mu\text{M} \cdot \text{s})^{-1}, k_b : 10\text{s}^{-1}$ | |
| $F + G \rightleftharpoons H$ | $k_f : 10(\mu\text{M} \cdot \text{s})^{-1}, k_b : 1\text{s}^{-1}$ | |
| $H + I \rightleftharpoons J$ | $k_f : 1(\mu\text{M} \cdot \text{s})^{-1}, k_b : 1\text{s}^{-1}$ | |

Simulation results of the simple model are summarized in Figure 8. Both STEPS 3 and STEPS 4 implementations demonstrate a steady decrease of simulation time early on until 2^6^ = 64 cores, and maintain roughly the same time cost for the rest of the configurations. The memory footprint improvement from STEPS 4 is significant. In the baseline simulations, STEPS 4 consumes 45.6MB of memory per core, about 60% of the required memory for STEPS 3. When simulating the model with thousands of cores, the memory consumption of STEPS 4 further decreases to about 4.5MB per core, 10% of the baseline simulation consumption, thanks to the completely distributed nature of the solution. While the memory footprint of STEPS 3 simulations also decreases with high core counts, the number stabilizes at 16MB, 2.6 times more than STEPS 4 requires. The strong scaling speedup for both STEPS 3 and STEPS 4 in Figure 8C suggests that the STEPS 4 achieves close-to-ideal speedup until 2^6^ = 64 cores, reflecting the time cost result in Figure 8A. In fact, the SSA component further maintains a linear speedup until 2^9^ = 512 cores according to the component scalability analysis in Figure 8D. However, due to the high scalability, its proportion in the overall time cost reduces significantly in high core count simulations. For these simulations, the diffusion operator and other background maintenance routines become the two major proportions of the simulation time cost.

**Figure 8:**
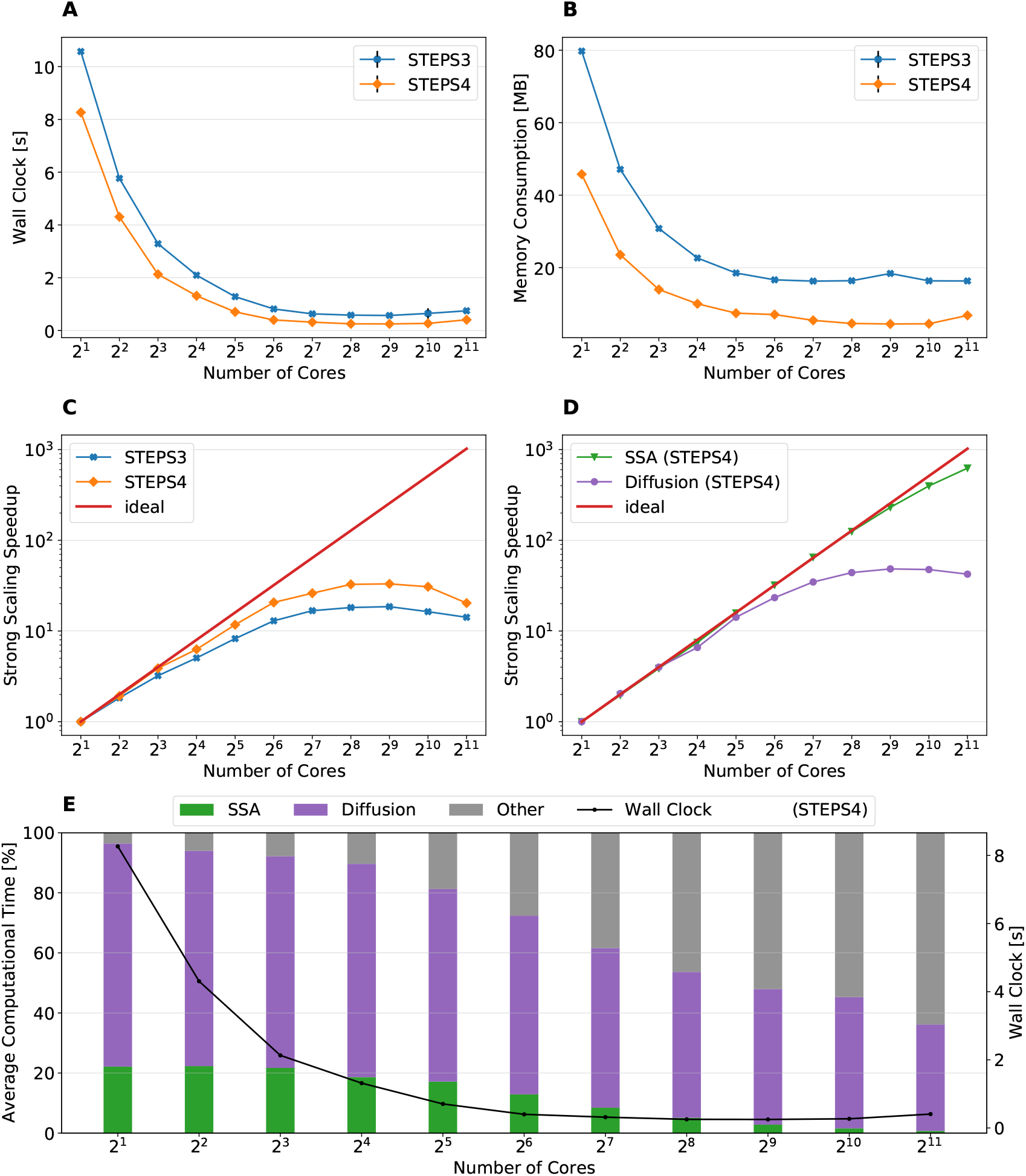
The performance results and scalability of the simple model. **(A)** Both STEPS 3 and STEPS 4 implementations demonstrate a steady decrease of time cost early on, then maintain similar time cost beyond 2^6^ = 64 cores. **(B)** STEPS 4 consumes significantly less per-core memory than STEPS 3, ranging from 60% in the baseline simulation, to approximately 30% in high core count simulations. **(C)** STEPS 4 achieves close-to-ideal speedup until 2^6^ = 64 cores, but has poor scalability afterward. Similar but slightly worse scalability can be observed for STEPS 3. **(D)** Component scalability analysis of STEPS 4 suggests a linear speedup of the SSA operator. **(E)** Component proportion analysis of STEPS 4. Because of the highly scalable SSA operator, the time cost of simulations with high core counts is dominated by the diffusion operator and other non-scalable bookkeeping routines, leading to the poor overall scalability in high core count scenarios.

#### 3.2.3 The Calcium Burst Background Model

We extend our investigation on the reaction-diffusion component with the the calcium burst background model with complex Purkinje cell morphology as described in section 3.1.3. The voltage component of this model is removed and only background channel reaction and diffusion is simulated. In total, the model consists of 15 molecule species, 8 of which are diffusive, and 22 reactions. The simulated mesh consists of 853,193 tetrahedrons. To eliminate any difference caused by partitioning, we pre-partition the mesh in Gmsh then import the partitioned mesh to the simulations, therefore the partitioning is always the same for each benchmark configuration. We start the simulation series from 2^5^ = 32 cores within a single node, then double the core count each time until the maximum of 512 nodes with 2^14^ = 16384 cores is reached.

Figure 9 presents the key results of the simulation series. In general, STEPS 4 performs slightly worse than STEPS 3 in low core count configurations, but eventually achieves similar performance as the core count increases. This is because currently STEPS 4 implements the widely accepted Gibson and Bruck (Gibson and Bruck, 2000) next reaction method as the default SSA operator. This method provides logarithmic computational complexity with simple data structures that we find suitable for the distributed solution. On the other hand, STEPS 3 inherits the serial implementation of the Composition and Rejection method (Slepoy et al., 2008), which requires a more complex data structure but takes advantage of its constant time complexity, particularly when dealing with large number of reactions in low core count simulations. It is worth noting that the compartmental design in STEPS 4 supports multiple operator implementations, therefore more efficient operators can be easily integrated to the solution in the future.

**Figure 9:**
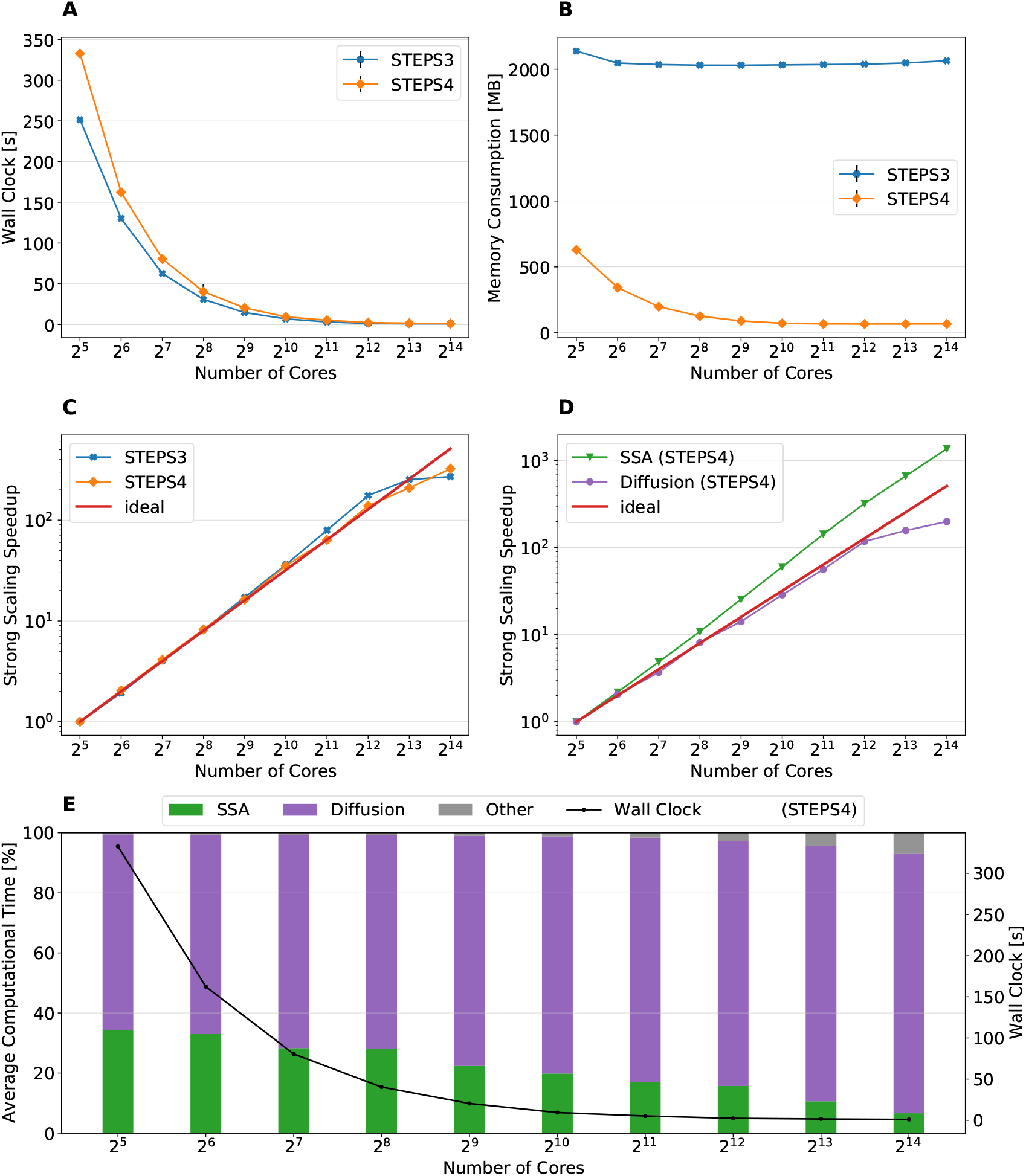
The performance results and scalability of the calcium burst background model. **(A)** Steady decrease of simulation time cost can be observed in both STEPS 4 and STEPS 3 simulations. STEPS 4 performs slightly worse than STEPS 3 in low core count simulations, both eventually achieves similar performance as core count increases. **(B)** The memory footprint of STEPS 4 is superior comparing to the STEPS 3 counterparts, requiring about 630MB for 2^5^ = 32 core simulations, and 67MB for 2^10^ = 1024 core and above simulations. STEPS 3 consumes more than 2GB of memory per core for the whole series. **(C)** Both STEPS 4 and STEPS 3 demonstrates linear to super-linear scaling speedup. **(D)** Component scalability analysis of STEPS 4 suggests that the diffusion operator in STEPS 4 exhibits linear speedup until 2^10^ = 1024 cores, while the SSA operator shows a remarkable super-linear speedup throughout the series. **(E)** Component proportion analysis of STEPS 4. The diffusion operator is the dominating component, taking from 65% to 95% of the overall computational time.

Dramatic improvement on memory consumption can be observed for STEPS 4 in Figure 9B. All STEPS 3 simulations require no less than 2GB of memory per core; on the other hand, the highest per-core memory footprint for STEPS 4 is about 630 MB with 2^5^ = 32 cores, and drops down to about 67MB with 2^10^ = 1024 cores and above, roughly 3% of what is required by STEPS 3.

Both STEPS 4 and STEPS 3 demonstrate linear to super-linear speedup until 2^12^ = 4096 cores in Figure 9C. Component scaling analysis in Figure 9D suggests that both the SSA and the diffusion operators contribute to this result. The diffusion operator maintains close-to-linear speedup until 2^12^ = 4096, while the SSA operator demonstrates super-linear speedup throughout the series. We investigated this scaling behavior and further profiling on the SSA operator indicates that the super-linear speedup mainly comes from the update routine of the operator, including the propensity calculations and the priority queue updates. This suggests that the improvement on memory caching may play an important role here.

The diffusion operator is the dominating component in this series, as shown in Figure 9E. Its proportion in the overall time cost increases from 65% to 95%. The proportion of other non-scaling routines also rises but is still less than 10% with the maximum 2^14^ = 16384 cores. Overall the performance profile of the background model is very similar to the simple model profile. This is not surprising as they both involve the same operators but the background model has more tetrahedrons and reactions per core compared to the simple model.

#### 3.2.4 Complete Calcium Burst Model

The complete calcium burst model as described in section 3.1.3 extends the background model by coupling molecular reaction-diffusion updates with voltage-dependent channel activation as well as membrane potential changes. Different channel density parameters are assigned to the smooth and the spiny sections of the mesh to approximate the effect caused by regional spine density difference. The model consists of 15 regular species, 8 of which are diffusive, 5 types of channels with in total 27 different channel states, 59 regular reactions and 16 voltage-dependent reactions. Compared to the previous two models, the complete model produces a simulation with extremely complex dynamics and imbalanced computational load, both spatially and temporally. We consider it as an excellent demonstration of STEPS 4 performance in realistic research projects.

Figure 10 summarizes the key results of the simulation series. While STEPS 4 performs slightly worse than STEPS 3 initially, it reaches similar performance with 2^9^ = 512 cores, and outperforms STEPS 3 for the rest of the series.

**Figure 10:**
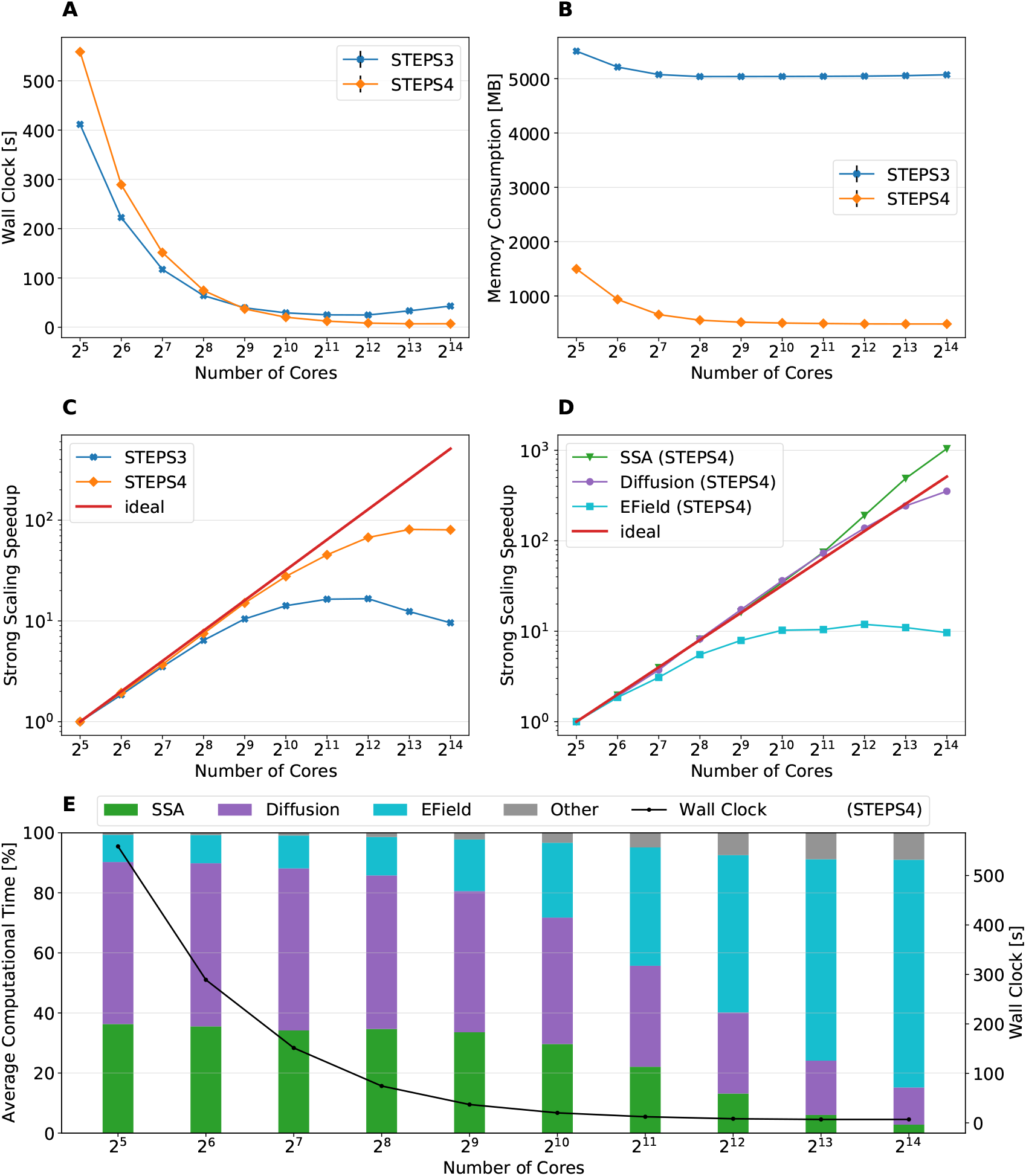
The performance results and scalability of the calcium burst complete model. **(A)** STEPS 4 performs slightly worse than STEPS 3 in low core count simulations, but reaches similar performance with 2^9^ = 512 cores, and outperforms STEPS 3 afterward. **(B)** STEPS 4 requires about 1.5GB for the 2^5^ = 32 core baseline simulations. Its memory footprint quickly decreases to 500MB for 2^9^ = 512 cores and above. STEPS 3 consumes more than 5GB of memory per core for the whole series. **(C)** STEPS 4 achieves a better scalability comparing to STEPS 3. **(D)** Component scalability analysis of STEPS 4. The SSA operator shows a super-linear speedup throughout the series. The diffusion operator also exhibits linear speedup until 2^13^ = 8192 cores. However, the EField operator shows limited scalability with maximum 10x speedup with 2^10^ = 1024 cores and above. **(E)** Component proportion analysis of STEPS 4. The EField operator progressively dominates the computational time, from 10% in the baseline simulations to 76% in the 2^14^ = 16384 core simulations, due to its limited scalability comparing to other operator components.

As expected, STEPS 4 continues its advantage on per-core memory footprint management, starting from 1.5GB for 2^5^ = 32 core simulations, to approximately 500MB for 2^9^ = 512 cores and above. The minimum memory requirement for STEPS 3 is 5GB, 10 times what is needed with STEPS 4. While the BB5 cluster has high memory capacity per compute node and is able to provide 12GB of memory per core for simulations (given the 32 active processes per node), many HPC clusters commonly have the memory capacity restriction of about 4GB per core (Zivanovic et al., 2017), therefore only STEPS 4 simulations may be run on those clusters.

Overall, STEPS 4 achieves a better scalability compared to STEPS 3, thanks to the linear speedup from the diffusion operator, and the super-linear speedup from the SSA operator. However, the EField operator has limited scalability, reaching maximum 10x speedup relative to the baseline. This results in a great increase of EField operator time cost in proportion to the total computation time, from 10% in the baseline simulations to 76% in the 2^14^ = 16384 core simulations, making it the major performance bottleneck of the series, as shown in Figure 10E.

#### 3.2.5 Memory Footprint with Refined Mesh

As shown in the above results, the significantly reduced memory footprint is one of the major advantages of STEPS 4. To further investigate the memory consumption difference between STEPS 4 and STEPS 3, we refine the Purkinje cell mesh and rerun both the calcium burst background model and the complete calcium burst model with the new mesh. The refined mesh consists of 3,176,770 tetrahedrons. For simplicity, we name the original mesh as the “1M” mesh, and the refined mesh as the “3M” mesh accordingly. As the 3M simulations exhibit similar performance profiles as the 1M versions, we skip the performance analysis and focus on the memory footprints of the simulations. Performance and scalability results of the 3M simulations can be found in the Supplementary Material Section 1.2.

Figure 11 provides an overall view of the results. For all simulation series, the baseline configuration has the highest memory footprint, then progressively reduces to a consistent minimum. We hereby use the minimum memory consumption from each series for comparison. Figure 11A presents the memory consumption of the background model, for both STEPS 3 and STEP4, and for both 1M and 3M meshes. For the 1M mesh simulations, STEPS 4 requires 67MB memory per core, while STEPS 3 requires approximate 2GB, 30x of the STEPS 4 requirement. For the 3M mesh simulations, 200MB memory per core is required by STEPS 4, while 6.6GB is required by STEPS 3, about 33x of the STEPS 4 requirement. Results of the complete model are shown in Figure 11B. For the 1M series, STEPS 4 requires about 500MB of memory, while STEPS 3 requires approximate 5.1GB, resulting in a 10x difference. For the 3M series, the memory footprint of STEPS 4 increases to 770MB. We are unable to simulate the 3M complete model with STEPS 3 as the 12GB per core memory capacity remains inadequate.

**Figure 11:**
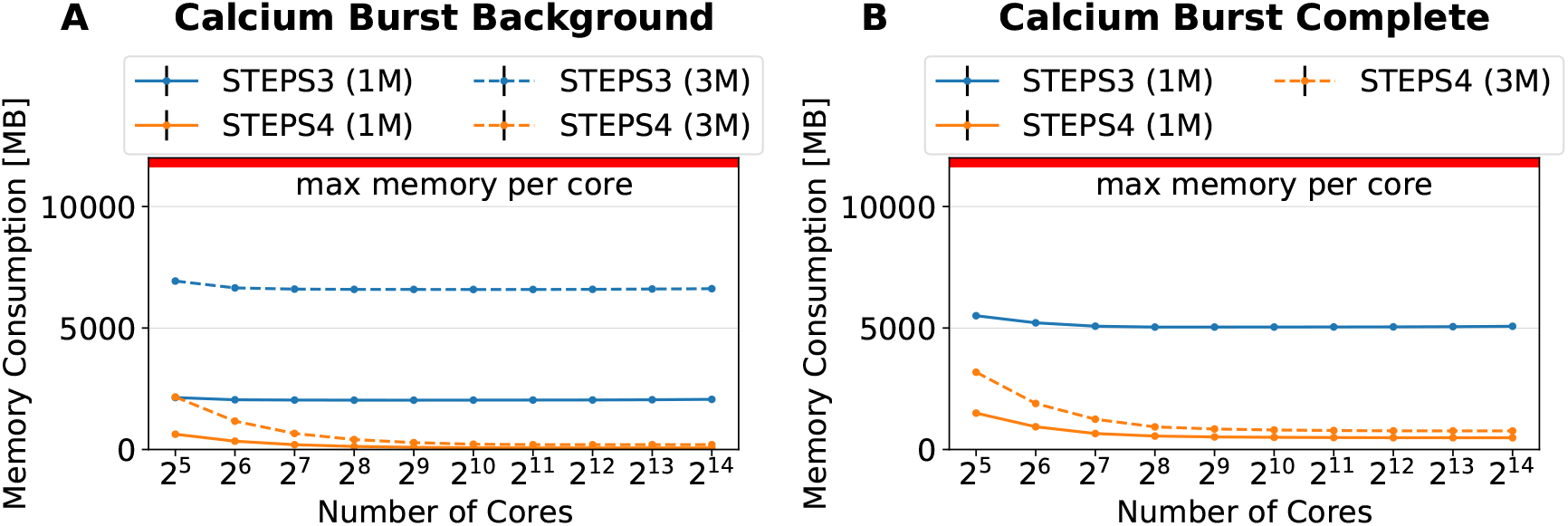
Memory footprints of the calcium burst background and complete model simulations with 1M and 3M meshes. **(A)** Results of the background model simulations. For the 1M series, STEPS 4 requires minimum 67MB per core, while STEPS 3 requires minimum 2GB. For the 3M series, STEPS 4 requires minimum 200MB, and STEPS 3 requires minimum 6.6GB. **(B)** Results of the complete model simulations. For the 1M series, STEPS 4 requires minimum 500MB per core, while STEPS 3 requires minimum 5.1GB. For the 3M series, STEPS 4 requires minimum 770MB. The 12GB per core memory capacity of the cluster is inadequate for the 3M complete model simulations with STEPS 3.

#### 3.2.6 Single Node Roofline Analysis of STEPS 4

In general, STEPS 4 demonstrates similar or better performance compared to STEPS 3 in high core count simulations, but has lower performance in small core count simulations. As discussed previously, one of the reasons is the different SSA operator implementations but other factors may also be involved. As the performance with small core count simulations is also important for STEPS 4 usage, a detailed performance analysis of current simulations is necessary to determine the direction of future optimizations. We choose the complete model as the profiling target since all major operators are included in the simulation. Note that in low core count configuration, the SSA and the diffusion operators are the dominating components in the simulation, thus they are the main focus of the analysis here. This is different from the optimization of high core count simulations, where the EField operator dominates the computation.

The analysis is based on the the Roofline model (Williams et al., 2009), evaluating the scaling trajectory (Ibrahim et al., 2018) of the most computationally expensive routines, in our case, the SSA reaction operator, the Diffusion operator, and the EField operator. The Roofline model is one of the simplest tools to apply hardware/software co-design, enabling investigation on the interaction between hardware characteristics like memory bandwidth and peak performance, and the software characteristics such as memory locality and arithmetic intensity. Thus it provides essential information on whether the investigated components are memory bandwidth or compute bound, and consequently vital suggestions on optimization strategies.

The Roofline model shown in Fig. 12 for the Cascade Lake node on BB5 is constructed from a measured memory bandwidth (≈ 197*GB/s*) and a measured peak core performance (≈ 78*Gflop/s*, where *flop* stands for floating-point operations). Both metrics are measured with the likwid-bench utility (Treibig et al., 2010). In the Roofline graph, the *x*-axis is the arithmetic (or computational) intensity (AI), computed as the ratio of floating point operations to transferred bytes from the main memory (DRAM traffic), and the *y*-axis is the observed performance. To obtain a scaling trajectory (Ibrahim et al., 2018) the measures are taken for varying core counts. Additionally, we run simulations with hyper-threading (2^6^ = 64 processes) in order to utilize maximum resources.

**Figure 12:**
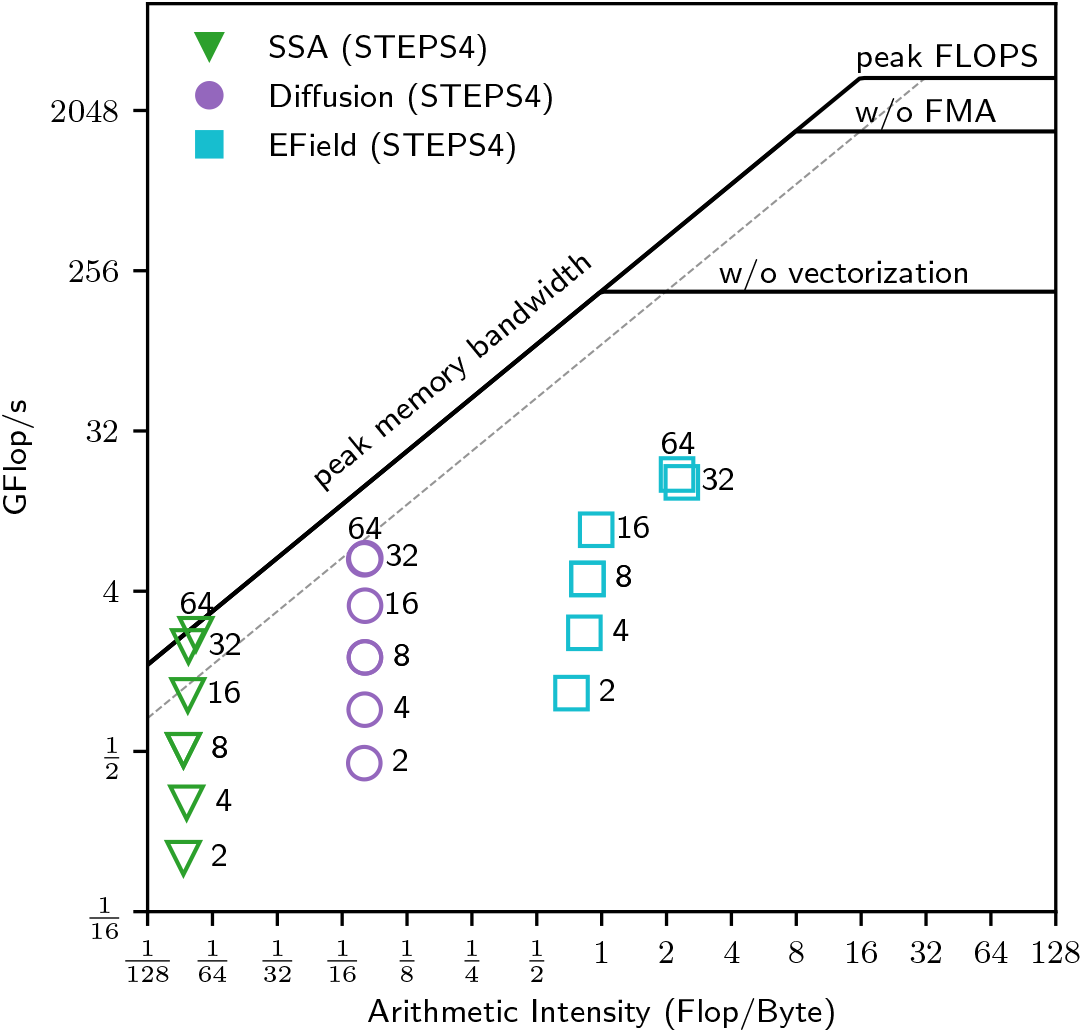
Roofline single-node scaling trajectories. The solid black lines are the full node hardware limits and the dashed gray line is the peak memory bandwidth for one socket. Each data point is labeled by the number of processes. All the computational kernels present a low arithmetic intensity mainly due to not ideal data locality (not optimal cache utilization). Nevertheless, the scaling is close to ideal (especially for the SSA and Diffusion operators) given that the doubling of concurrency leads to a corresponding Δ*y >* 0. Hyper-threading at 64 cores does not give any substantial performance increase.

For the measurements of the routine with LIKWID, a MPI synchronization barrier is added before and after each measured kernel. This is done to ensure that the measured metrics (e.g. hardware counters) indeed belong to the respective routines.

From Figure 12, it can be seen that all routines have a low arithmetic intensity. Each routine is represented by a different symbol and each data point is labeled by the number of processes. As described in Ibrahim et al. (2018), for ideal scaling, a doubling of concurrency corresponds to a change in Δ*y* (observed Flop/s) of ≈2 × without a corresponding change in Δ*x* (arithmetic intensity), a behavior observed in our experiments. The SSA kernel is the one with the lowest arithmetic intensity (AI) and it is well into the AI regime where we expect the kernel to be memory bound. The Diffusion kernel presents similar behavior but with higher arithmetic intensity. Both kernels reach a saturation point as they approach the peak memory bandwidth. This observation suggests that there would be little to no gain to be had by vectorizing these kernels, instead possible improvements would have to come from algorithmic changes and/or cache blocking strategies in order to either increase the arithmetic intensity or fit the working memory set into the last level cache (LLC). For the EField kernel, we observe both Δ*y* > 0 and Δ*x* > 0 as we perform the strong scaling. This transition indicates that the number of floating-point operations has remained constant, so data movement must have decreased (Ibrahim et al., 2018). Finally, for all the computational kernels hyper-threading does not lead to any substantial performance increase.

To reach the maximum performance of a compute node, we need to efficiently utilize the cache memory hierarchy. In the Roofline graph, the higher the cache efficiency the higher the computational intensity. In our case, the low arithmetic intensity could be explained by the use of data structures that do not favor data locality (e.g. maps/dictionaries over vectors). Thus, a substantial improvement in the computational intensity of STEPS 4 can be achieved by favoring data locality and thus higher cache utilization. Towards this direction, we already mentioned in 2.2 the `flat-multimap` container, which we plan on using extensively throughout STEPS 4 codebase.

## 4 Discussion

In the present manuscript we achieved several major goals:

- We developed a more user-friendly modeling environment by introducing the new API2 system to STEPS 4 development.
- We modernized the existing code base of the entire framework adopting modern programming standards and practices such as C++17 and continuous integration. Particular care was posed on safety features such as strong type indexing. These improvements provided a solid modern foundation for STEPS 4 development.
- A distributed solution that addressed the bottlenecks of STEPS 3.

STEPS 4 achieves similar performance and scalability as STEPS 3 while dramatically reducing the memory footprint. This is a key feature for future realistic modeling using STEPS. The Purkinje dendrite morphology simulated in the calcium burst model was reconstructed from light microscopic imaging. The spines were ignored and only the skeleton of the dendrite was preserved. It is possible to reconstruct a highly realistic Purkinje neuron containing all visible spines from high resolution electron microscopic imaging, however, the mesh generated from such morphology is expected to have 10 to 100 times more tetrahedrons than the one used in current simulations. Such large models are completely out-of-reach for STEPS 3 since even the relatively small 3M calcium burst model already exceeds the 12GB per-core memory capacity on a state-of-the-art cluster like BB5. Conversely, STEPS 4 showed its potential on supporting super large scale simulations thanks to its distributed nature.

### 4.1 Limitations and solutions

STEPS 4 is not a complete replacement for STEPS 3. It is a highly specialized version of the operator-splitting solution specifically tailored for cluster-based, super-large scale simulations. Thus, we paid particular attention to performance optimizations whilst maintaining accuracy.

Even if STEPS 4 covers most features available in STEPS 3, some remain missing. For instance, the diffusion of species on surfaces (i.e. between patch triangles), and the associated surface diffusion boundaries, are not yet available. Patches between compartments are in principle supported but meshes have to be partitioned in such a way that tetrahedrons on both sides of patch triangles are owned by the same process. Differently from STEPS 3 where automatic partitioning adjustment is provided, this constraint is currently unenforced by Omega_h and STEPS 4. It relies on the modelers to generate suitable partitioned meshes for their simulations. We plan to support automatic partitioning adjustment with constraints in STEPS 4, however, this requires further collaboration with the Gmsh and Omega_h developers as these libraries need further development to support such functionality. Finally, some auxiliary features in STEPS 3 such as the Region of Interest (ROI) functionality and visualization are not yet supported as implementations of new STEPS modeling toolkits are required to adapt the new distributed mesh formats and protocols.

### 4.2 Potential enhancements for STEPS 4

The distributed mesh backend of STEPS 4, Omega_h, not only supports traditional MPI based distributed-memory parallelism, but also shared-memory parallelism through OpenMP, and GPU parallelization via the CUDA framework. It also provides unique features such as mesh adaptation suitable for GPUs using flat array data structures and bulk transformations. These advanced features are currently not utilized in STEPS 4 as it relies solely on CPU based MPI parallelism. With the importance of GPU based fat compute nodes in modern HPC clusters, such features will play important role when STEPS 4 is transitioned to other parallelism schemes.

In addition, the scalability analysis in Section 3.2 suggests two major axes for future development. On one hand, the EField operator is shown to be the major bottleneck in high core count simulations due to its poor scalability. On the other hand, the Roofline analysis shows low computational intensity for all major kernels (SSA, Diffusion, EField). This behavior points to unsatisfactory use of cache memory, mainly caused by containers/data structures with poor data locality. A more extensive use of the `flat-multimap` could greatly improve cache utilization and increase the arithmetic intensity of these computational kernels.

### 4.3 Other current developments and future directions

#### 4.3.1 Vesicle modeling

Currently STEPS, as also all the SSA methods in general, models molecules as points that do not occupy a significant volume of the space in which they reside. This is an obvious limitation if one wants to model certain types of structures in the cell such as vesicles. Vesicles are relatively large structures (*∼* 40nm diameter in the case of synaptic vesicles for example) that play many important roles in biology, and their complex structure and diverse functionality mean they cannot be realistically simulated by the point-molecule approach. Vesicles undergo processes such as endocytosis and exocytosis, interact with cytosolic and surface-bound molecules, and can be spatially organized into clusters such as in the presynaptic readily-retrievable pool. In an upcoming release, STEPS aims to support all of these features in an initial parallel implementation.

While the vesicle modeling development has been a separate project from STEPS 4, one tantalizing prospect is to marry many of the novel features of STEPS 4 with the vesicle modeling to allow bigger, more detailed simulations that can be run for longer biological times.

#### 4.3.2 Coupling of STEPS with other simulator software

Recently, the STEPS team at OIST and the Blue Brain Project (BBP) at EPFL have been collaborating in software development and scientific research such as simulation support for multi-scale modeling. As part of the BBP mission to create a large scale reconstruction of brain tissue, a multi-scale approach for simulation is deemed necessary to capture elements at various temporal and spatial scales. One time scale for rapidly changing neuron voltages, a different, slower time scale for changing ion concentrations. Likewise, neuron morphologies can be distributed among computing ranks irrespective of geometric boundaries whereas bulk ion concentrations and metabolism use a coarse grain division of the spatial scale. For this purpose different simulators are used to leverage their specialized capabilities. NEURON (Carnevale and Hines, 2009) is used to solve relevant equations for membrane voltage and communication between neurons in addition to calcium in astrocyte morphologies. Meanwhile STEPS is employed to compute concentrations of diffusing ions in the extracellular space. A more memory efficient STEPS better enables sharing of computing resources between the two simulators.

## 5 Conclusion

The STEPS 4.0 project development reported in this article addresses several issues in previous STEPS releases, improving the user modeling experience, as well as modernizing the existing code base in order to aid future developments. The main contribution of this research is a new parallel stochastic reaction-diffusion solver supported by a sophisticated distributed mesh library. While maintaining similar performance and scalability, the new solver dramatically reduces the memory footprint of simulations, resolving the major bottleneck in previous solutions. This breakthrough is essential to future neuroscience research as it enables super-large scale molecular reaction-diffusion simulations with biologically realistic models. Into the future, the STEPS project aims to continue to innovate, bringing new features, realism and detail to molecular neuronal models. Such efforts cater to researchers as they ask new questions and investigate new problems through computer simulation to advance our understanding of the brain and other biological systems.

## Author contributions

EDS and FS conceptualized and led this study. TC and WC led the overall software development of STEPS 4. SM and TC added in Omega_h the support to Gmsh file format 4 and features required in STEPS 4. BDM, SM, TC and WC contributed to the Zee library development and evaluations. AC, BDM, CK, GC, JL, SM, TC and WC contributed to the software development of STEPS 4. AC, CK, IH, JL, TC and WC contributed to the pre-release testing, debugging and optimization of STEPS 4. JL contributed to the API2 development for STEPS 4 and model conversions from STEPS 3 to STEPS 4. AC and IH designed and conducted the validation benchmarks. CK, GC and WC designed and conducted the performance benchmarks. NC checked statistical soundness of the tests and contributed in CI. OA, JK and PK contributed to technical discussions and supervised the BBP team. WC coordinated the writing of the manuscript. AC, BDM, CK, EDS, FS, GC, IH, JK, JL, NC, OA, PK, SM, TC and WC contributed to the manuscript. All authors gave feedback on the manuscript.

## Data Availability Statement

The STEPS simulator is available at http://steps.sourceforge.net/. Models for validation and performance investigation presented in this publication are available at https://github.com/CNS-OIST/STEPS4ModelRelease/.

## Footnotes

1 https://github.com/CNS-OIST/STEPS_PolyhedronROI

2 http://steps.sourceforge.net/manual/manual_index.html

3 https://github.com/CNS-OIST/STEPS_Validation

4 http://neuromorpho.org/neuron_info.jsp?neuron_name=10-2012-02-09-001

## References

Abhyankar, S., Brown, J., Constantinescu, E. M., Ghosh, D., Smith, B. F., and Zhang, H. (2018). Petsc/ts: A modern scalable ode/dae solver library. arXiv preprint 1806.01437

Anwar, H., Hepburn, I., Nedelescu, H., Chen, W., and De Schutter, E. (2013). Stochastic calcium mechanisms cause dendritic calcium spike variability. Journal of Neuroscience 33, 15848–15867. doi:10.1523/JNEUROSCI.1722-13.2013

Anwar, H., Roome, C. J., Nedelescu, H., Chen, W., Kuhn, B., and De Schutter, E. (2014). Dendritic diameters affect the spatial variability of intracellular calcium dynamics in computer models. Frontiers in Cellular Neuroscience 8, 168

Ascoli, G. A., Donohue, D. E., and Halavi, M. (2007). NeuroMorpho.Org: a central resource for neuronal morphologies. J Neurosci 27, 9247–9251

Bhalla, U., Bilitch, D., and Bower, J. (1992). Rallpacks: A set of benchmarks for neuronal simulators. Trends in neurosciences 15, 453–8. doi:10.1016/0166-2236(92)90009-W

Boehme, D., Gamblin, T., Beckingsale, D., Bremer, P.-T., Gimenez, A., LeGendre, M., et al. (2016). Caliper: performance introspection for HPC software stacks. In SC’16: Proceedings of the International Conference for High Performance Computing, Networking, Storage and Analysis (IEEE), 550–560

Carnevale, N. T. and Hines, M. L. (2009). The NEURON Book (USA: Cambridge University Press), 1st edn.

Chen, W. and De Schutter, E. (2017). Parallel STEPS: Large scale stochastic spatial reaction-diffusion simulation with high performance computers. Frontiers in Neuroinformatics 11, 13. doi:10.3389/fninf.2017.00013

Chen, W., Hepburn, I., Martyushev, A., and De Schutter, E. (in press). Modeling neurons in 3d at the nanoscale. In Computational Neuroscience Approaches to Cells and Circuits, ed. M. Guigliano (Springer Nature)

Chylek, L. A., Stites, E. C., Posner, R. G., and Hlavacek, W. S. (2013). Innovations of the rule-based modeling approach. In Systems Biology (Springer). 273–300

Denizot, A., Arizono, M., Nägerl, U. V., Soula, H., and Berry, H. (2019). Simulation of calcium signaling in fine astrocytic processes: Effect of spatial properties on spontaneous activity. PLoS computational biology 15, e1006795–e1006795. doi:10.1371/journal.pcbi.1006795.31425510[pmid]

Gamblin, T., LeGendre, M., Collette, M. R., Lee, G. L., Moody, A., de Supinski, B. R., et al. (2015). The Spack Package Manager: Bringing Order to HPC Software Chaos (Austin, Texas, USA), Supercomputing 2015 (SC’15). doi:10.1145/2807591.2807623.LLNL-CONF-669890

Geuzaine, C. and Remacle, J.-F. (2009). Gmsh: A 3-D finite element mesh generator with built-in pre- and post-processing facilities. International Journal for Numerical Methods in Engineering 79, 1309–1331. doi:10.1002/nme.2579

Gibson, M. A. and Bruck, J. (2000). Efficient exact stochastic simulation of chemical systems with many species and many channels. The Journal of Physical Chemistry A 9, 104

Gillespie, D. T. (1977). Exact stochastic simulation of coupled chemical reactions. The Journal of Physical Chemistry 81, 2340–2361. doi:10.1021/j100540a008

Hepburn, I., Cannon, R., and De Schutter, E. (2013). Efficient calculation of the quasi-static electrical potential on a tetrahedral mesh and its implementation in steps. Frontiers in Computational Neuroscience 7, 129. doi: 10.3389/fncom.2013.00129

Hepburn, I., Chen, W., and De Schutter, E. (2016). Accurate reaction-diffusion operator splitting on tetrahedral meshes for parallel stochastic molecular simulations. The Journal of Chemical Physics 145, 054118. doi:10.1063/1.4960034

Hepburn, I., Chen, W., Wils, S., and De Schutter, E. (2012). STEPS: efficient simulation of stochastic reaction–diffusion models in realistic morphologies. BMC Systems Biology 6, 36. doi:10.1186/1752-0509-6-36

Hodgkin, A. L. and Huxley, A. F. (1952). A quantitative description of membrane current and its application to conduction and excitation in nerve. J Physiol 117, 500–544

Ibanez, D. and Roberts, N. (2018). Omega_h. [Software] https://github.com/sandialabs/omega_h. doi:10.11578/dc.20180301.2

Ibrahim, K., Williams, S., and Oliker, L. (2018). Roofline scaling trajectories: a method for parallel application and architectural performance analysis. In 2018 International Conference on High Performance Computing & Simulation (HPCS) (IEEE), 350–358

Insel, T. R., Landis, S. C., and Collins, F. S. (2013). The nih brain initiative. Science 340, 687–688. doi:10.1126/science.1239276

Markram, H., Meier, K., Lippert, T., Grillner, S., Frackowiak, R., Dehaene, S., et al. (2011). Introducing the human brain project. Procedia Computer Science 7, 39–42. doi:https://doi.org/10.1016/j.procs.2011.12.015. Proceedings of the 2nd European Future Technologies Conference and Exhibition 2011 (FET 11)

Mohapatra, N., Tønnesen, J., Vlachos, A., Kuner, T., Deller, T., Nägerl, U. V., et al. (2016). Spines slow down dendritic chloride diffusion and affect short-term ionic plasticity of gabaergic inhibition. Scientific reports 6, 23196–23196. doi:10.1038/srep23196.26987404[pmid]

Rodola, G. (2020). psutil. https://github.com/giampaolo/psutil.v5.8.0

Schelker, M., Mair, C. M., Jolmes, F., Welke, R.-W., Klipp, E., Herrmann, A., et al. (2016). Viral rna degradation and diffusion act as a bottleneck for the influenza a virus infection efficiency. PLOS Computational Biology 12, 1–23. doi:10.1371/journal.pcbi.1005075

Slepoy, A., Thompson, A. P., and Plimpton, S. J. (2008). A constant-time kinetic Monte Carlo algorithm for simulation of large biochemical reaction networks. J Chem Phys 128, 205101

Sodani, A., Gramunt, R., Corbal, J., Kim, H.-S., Vinod, K., Chinthamani, S., et al. (2016). Knights landing: Second-generation intel xeon phi product. IEEE Micro 36, 34–46. doi:10.1109/MM.2016.25

Stillman, N. R., Balaz, I., Tsompanas, M.-A., Kovacevic, M., Azimi, S., Lafond, S., et al. (2021). Evolutionary computational platform for the automatic discovery of nanocarriers for cancer treatment. NPJ Computational Materials 7, 1–12

Treibig, J., Hager, G., and Wellein, G. (2010). LIKWID: A lightweight performance-oriented tool suite for x86 multicore environments. In Proceedings of PSTI2010, the First International Workshop on Parallel Software Tools and Tool Infrastructures (San Diego CA)

Trott, C. R., Lebrun-Grandié, D., Arndt, D., Ciesko, J., Dang, V., Ellingwood, N., et al. (2022). Kokkos 3: Programming model extensions for the exascale era. IEEE Transactions on Parallel and Distributed Systems 33, 805–817. doi:10.1109/TPDS.2021.3097283

Williams, S., Waterman, A., and Patterson, D. (2009). Roofline: An insightful visual performance model for floating-point programs and multicore. ACM Communications

Wils, S. and De Schutter, E. (2009). Steps: modeling and simulating complex reaction-diffusion systems with python. Frontiers in Neuroinformatics 3, 15. doi:10.3389/neuro.11.015.2009

Zamora Chimal, C. G. and De Schutter, E. (2018). Ca2+ requirements for long-term depression are frequency sensitive in purkinje cells. Frontiers in Molecular Neuroscience 11, 438. doi:10.3389/fnmol.2018.00438

Zivanovic, D., Pavlovic, M., Radulovic, M., Shin, H., Son, J., Mckee, S. A., et al. (2017). Main memory in hpc: Do we need more or could we live with less? ACM Trans. Archit. Code Optim. 14. doi:10.1145/3023362

