## supplementary material for "STEPS 4.0: Fast and memory-efficient molecular simulations of neurons at the nanoscale"

A PREPRINT

March 26, 2022

#### 1 Supplementary Data

##### 1.1 Validations

###### 1.1.1 Rallpack 1: Mesh refinement

We further examine the model convergence through mesh refining. Usually such studies start from a very coarse mesh and refine the solution through mesh splitting until results converge and residual errors plateau at numerical precision. However, the cable geometry is extreme: its length is 3-4 orders of magnitude bigger than its radius. This means that sensible mesh refinements split it only lengthwise. Since $V_{z_{min}}$ is the voltage trace where the current is injected and is dominated by the mesh size in the radial direction, it can be discarded in the current analysis. The focus is on $V_{z_{max}}$. Figure 1 presents the results in the typical log-log scale. As expected accuracy plateaus as it approaches numerical precision.

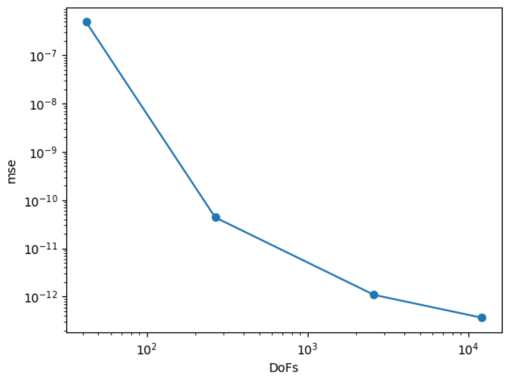


Figure 1: Convergence of the STEPS 4 trace $V_{z_{max}}$ to the analytical solution through mesh refinement in a log-log graph.

##### 1.1.2 Rallpack 3: Distributions, means and std. deviations

Here we present the distribution plots for the key features presented before. This is not meant to be proof that the two simulators produce the same results. It is to introduce the reader to the results with a general, qualitative overview.

Since peak heights and time stamps resemble bell-shaped curves, it is also interesting to compare means and standard deviations.

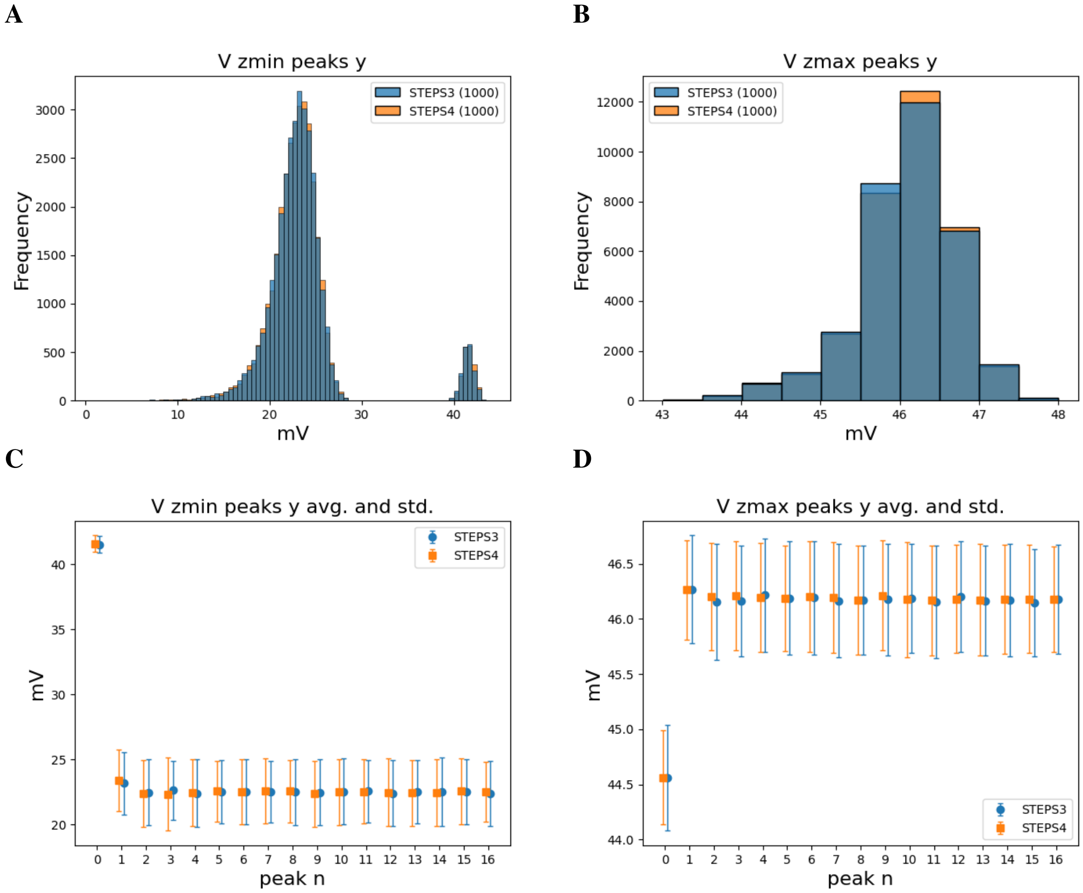


Figure 2: Distribution of peak heights for STEPS 3 and 4, mean and std. deviation. As mentioned before the plot for $z_{min}$ presents two different peaks because the first spike in the spike train is consistently higher. In addition we notice that there is much less variance at $z_{max}$ than $z_{min}$; sign that the current, flowing through the cable, averages out small imbalances introduced with the mesh. Qualitatively, STEPS 3 and 4 are in agreement.

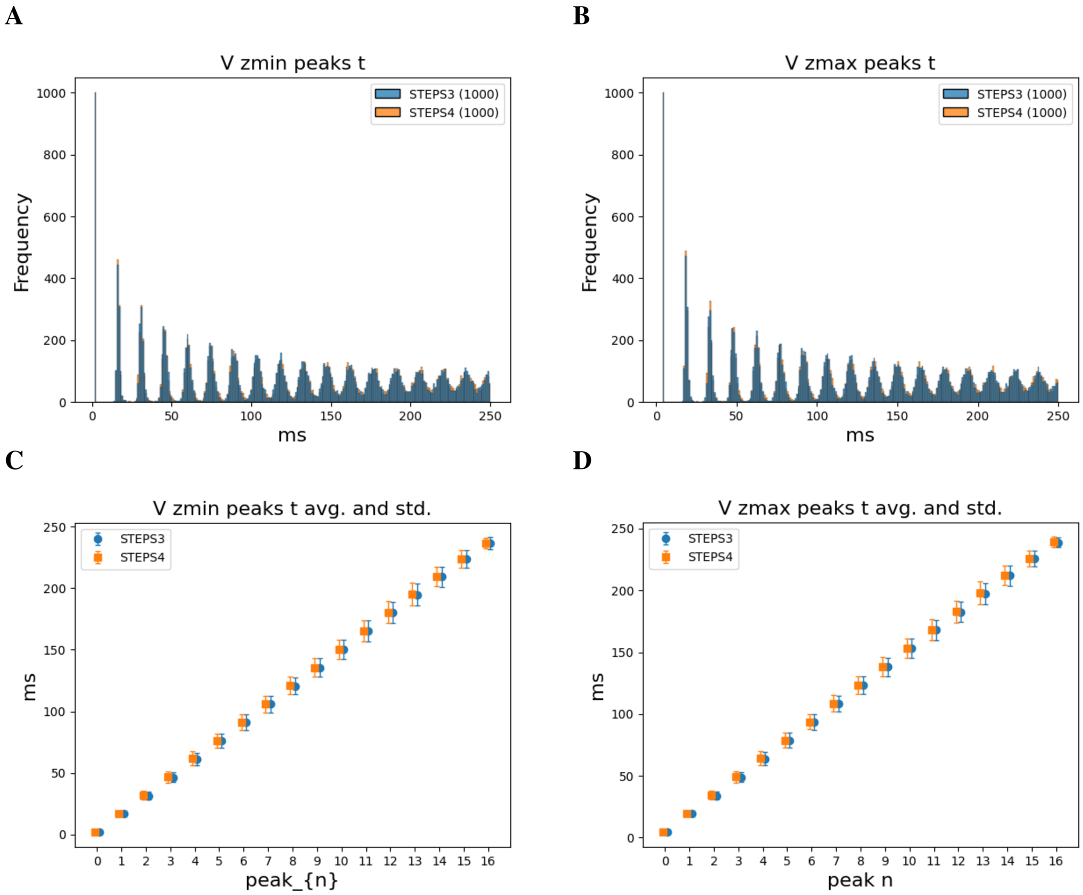


Figure 3: Distribution ((**A**) and (**B**)), mean and std. deviation ((**C**) and (**D**)) of peak time stamps for STEPS 3 and 4. The first peak is deterministic: appears always at the same time and the std. deviation is small. After, randomness enters the system and peaks tend to go out of phase, lowering peaks in the distribution analysis and widening the bases. Means and std. deviations climb as time passes. STEPS 3 and 4 results overlap.

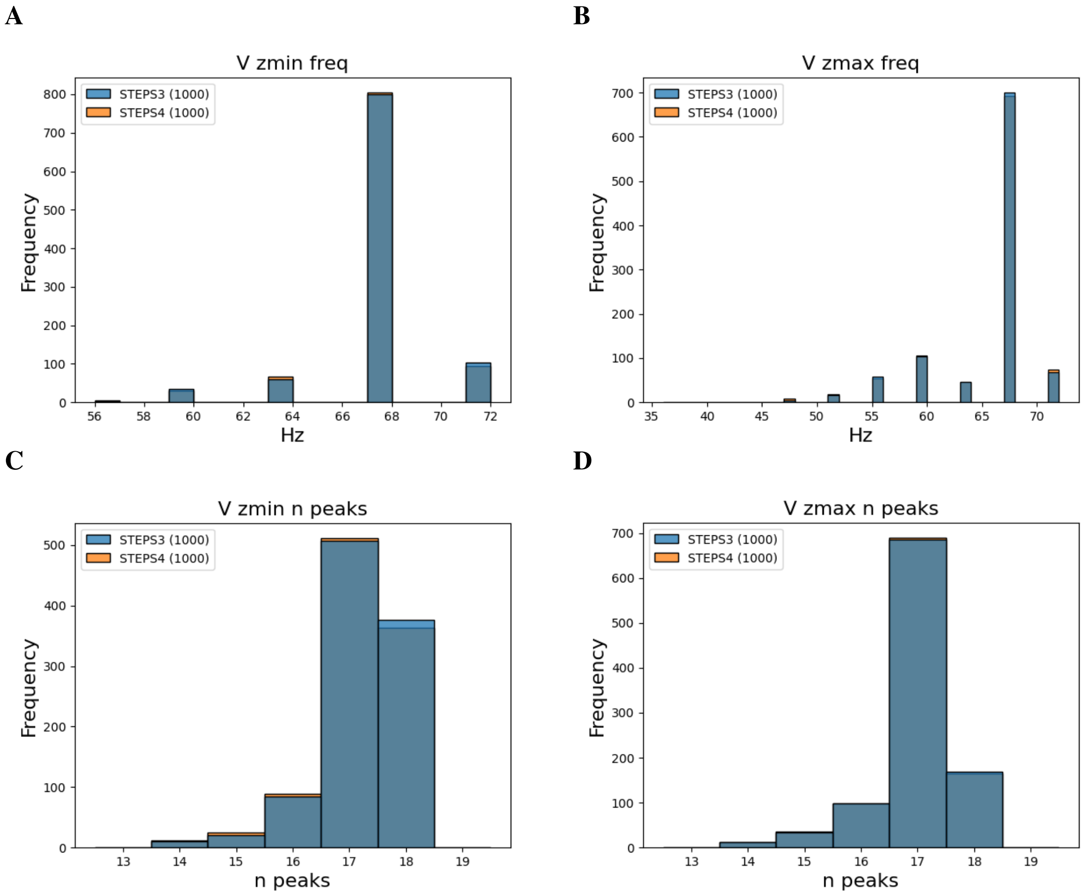


Figure 4: Distribution of the frequencies ((**A**) and (**B**)) and number of peaks ((**C**) and (**D**)) for STEPS 3 and 4. Usually in 0.25 s 17 peaks appear and the frequency is ~69 Hz. However, there are a few cases where a peak or two is skipped. There are also cases where the simulation is a little faster and an additional peak appears. In all the cases, STEPS 3 and 4 present the same behavior.

##### 1.2 Performance

We refined the Pukinje cell mesh and rerun both the calcium burst background model and the complete model. The refined mesh consists of 3,176,770 tetrahedrons and it is $\approx 3\times$ larger than the original one. Figure 5 presents the profiling of the calcium burst background model, and Figure 6 the corresponding metrics for the calcium burst complete model.

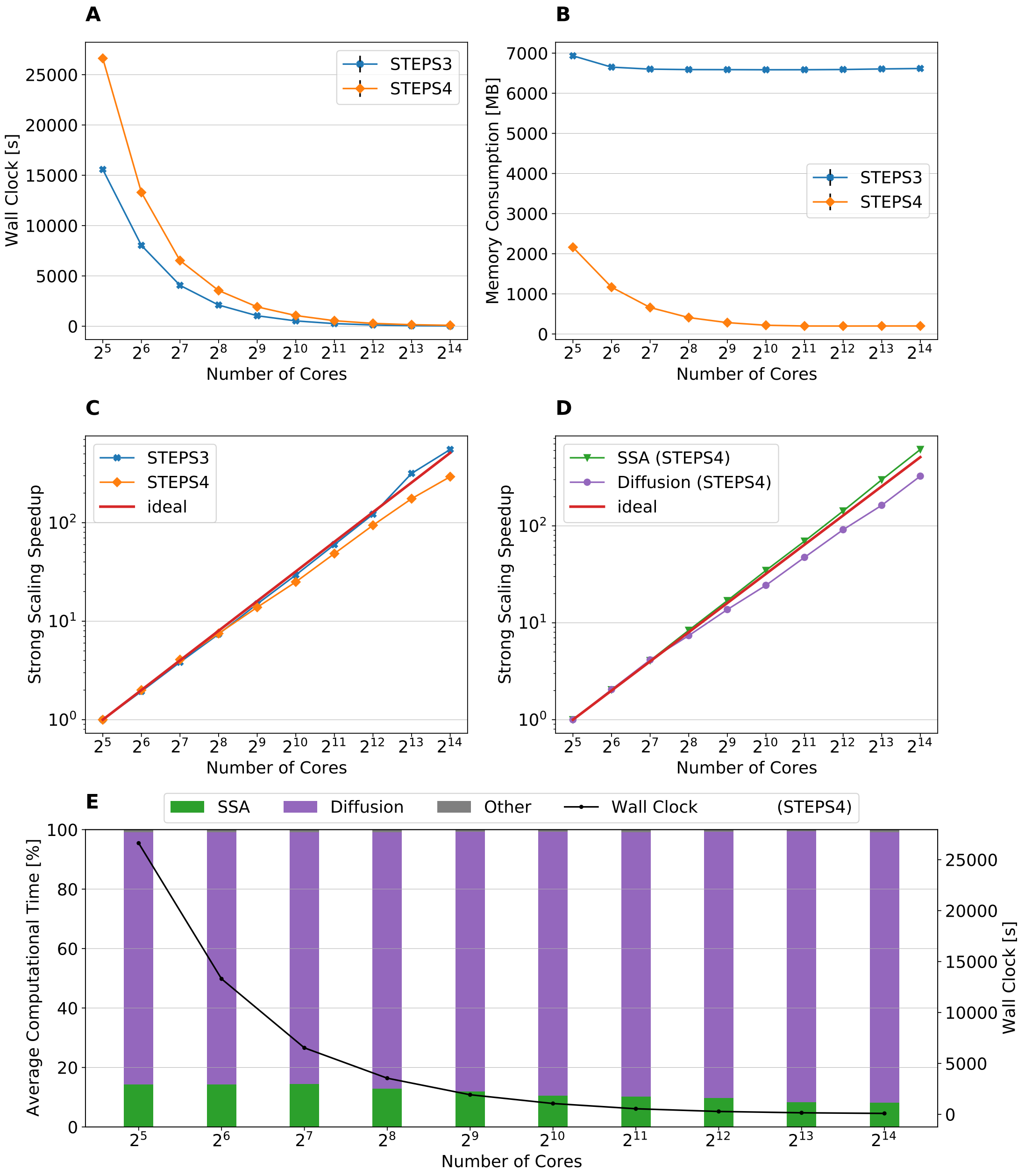


Figure 5: The performance results and scalability of the calcium burst background model for the refined mesh (denoted as "3M"). **(A)**: Steady decrease of simulation time cost can be observed in both STEPS 4 and STEPS 3 simulations. STEPS 4 performs slightly worse than STEPS 3 in low core count simulations, both eventually achieves similar performance as core count increases. **(B)**: The memory footprint of STEPS 4 is superior compared to the STEPS 3 counterparts, requiring about 2GB for $2^5$ core simulations, and 200MB for $2^{10}$ core and above. STEPS 3 consumes more than 6.5GB of memory per core for the whole series. **(C)**: Both STEPS 4 and STEPS 3 demonstrates linear to super-linear scaling speedup. **(D)**: The diffusion operator in STEPS 4 exhibits close to linear speedup until $2^9$ cores, while the SSA operator shows a remarkable super-linear speedup throughout the series. **(E)**: The diffusion operator is the dominating component, taking from 85% to 90% of the overall computational time.

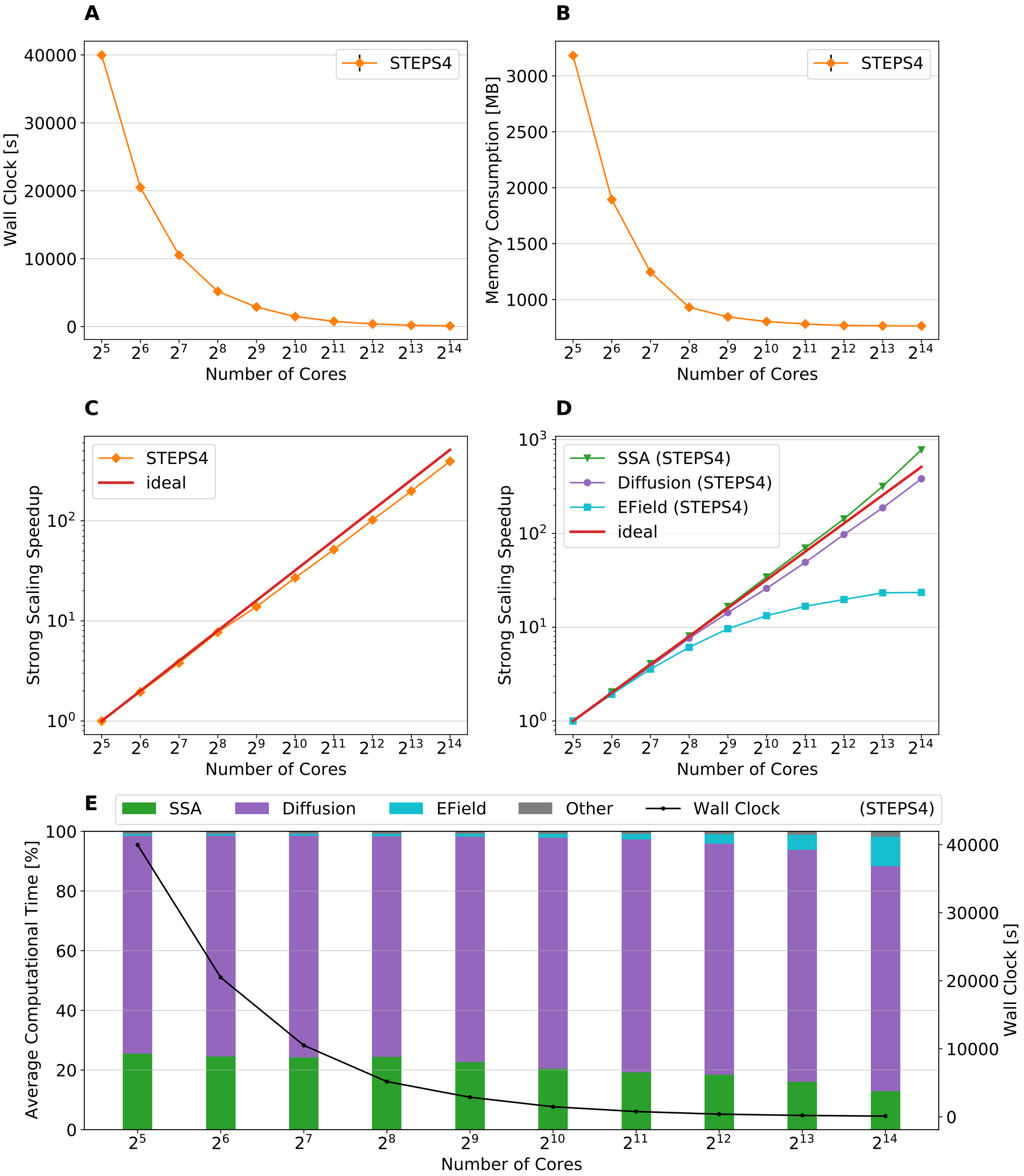


Figure 6: The performance results and scalability of the calcium burst complete model for the refined mesh (denoted as "3M"). We are unable to simulate the "3M" complete model with STEPS 3 as the 12GB per core memory quota is surpassed. **(A)**: STEPS 4 Wall Clock time. **(B)**: STEPS 4 memory consumption per core which is way below the 12GB per core memory quota that STEPS 3 exhausted. **(C)**: STEPS 4 achieves a close to linear speedup. **(D)**: The SSA operator shows a super-linear speedup throughout the series. The diffusion operator also exhibits close to linear speedup. However, the EField operator shows limited scalability. **(E)**: Average Computational Time per component.

#### 2 Supplementary Tables

Table 1: Examples of original (left column) and API2 (right column) usage for common parts of STEPS python scripts. These examples are extracts from larger python scripts and are not usable on their own. The new API has been designed to improve the readability of STEPS scripts. For example, chemical reactions are declared with a syntax that resembles chemical reaction notation and reactions in the same volume system are grouped together visually by using context managers (`with` statements). In addition to the increased readability, effort has also been put into reducing the amount of code that STEPS users need to write for common tasks like data saving. Data can be described with `ResultSelectors` and STEPS will take care of saving it automatically.

<table>
<tr><th></th><th>Original API</th><th>API2</th></tr>
<tr><td>Model declaration</td><td><pre>mdl = smodel.Model()
### Species declarations
SA = smodel.Spec('SA', mdl)
SB = smodel.Spec('SB', mdl)
vsys = smodel.Volsys('vsys', mdl)
### Reactions declarations
R1 = smodel.Reac('R1', vsys,
        lhs=[SA, SB], rhs=[SB, SB],
        kcst=K_f)
R2 = smodel.Reac('R2', vsys,
        lhs=[SB, SB], rhs=[SA, SB],
        kcst=K_b)</pre></td><td><pre>mdl = Model()
r = ReactionManager()
with mdl:
    # Species declarations
    SA, SB = Species.Create()
    vsys = VolumeSystem.Create()
    with vsys:
        # Reactions declarations
        SA + SB <r[1]> 2*SB
        r[1].K = K_f, K_b</pre></td></tr>
<tr><td>Data saving</td><td><pre>tpnts = numpy.arange(0, END_TIME, DT)
vSA = numpy.zeros(len(tpnts))
vSB = numpy.zeros(len(tpnts))

sim.reset()
### Set the initial conditions
sim.setCompCount('comp1', 'SA', nSA)
sim.setCompCount('comp1', 'SB', nSB)

for i in range(len(tpnts)):
    # Run the simulation
    sim.run(tpnts[i])
    # Save data
    vSA[i] = sim.getCompCount(
        'comp1', 'SA')
    vSB[i] = sim.getCompCount(
        'comp1', 'SB')</pre></td><td><pre># Declare the data to be saved
rs = ResultSelector(sim)
vals = rs.comp1.ALL(Species).Count
sim.toSave(vals, dt=DT)

sim.newRun()
### Set the initial conditions
sim.comp1.SA.Count = nSA
sim.comp1.SB.Count = nSB

### Run the whole simulation with
### automatic data saving
sim.run(END_TIME)</pre></td></tr>
</table>
